# Curved microtubule regions mark sites of lattice compaction in cells and neurons

**DOI:** 10.64898/2026.07.08.737175

**Authors:** Jaya Mishra, Polina Volos, Ke Wang, Barbara Birk, Laura Trefftz, Serapion Pyrpassopoulos, Nisha Mohd Rafiq

**Affiliations:** Interfaculty Institute of Biochemistry, University of Tübingen, Auf der Morgenstelle 34, 72076 Tübingen, Germany; Center for Plant Molecular Biology (ZMBP), University of Tübingen, Auf der Morgenstelle 32, 72076 Tübingen, Germany

**Keywords:** Tau, MAPs, C1-domain proteins, osmotic stress, iPSC-derived neurons

## Abstract

Angstrom-scale changes in microtubule (MT) lattice spacing regulate the selective recruitment of MT-associated proteins, yet how these structural states operate in cells remains poorly understood. Here, we show that MT lattice expansion, induced by protein-based expanders or microtubule-stabilizing agents such as Taxol and epothilone D, drives the relocalization of compact lattice–binding proteins, including tau, doublecortin (DCX), and the C1 domain–containing signaling protein GEF-H1, into highly curved MT-associated domains, whereas the compaction-inducing agent laulimalide suppresses this response. In contrast, the tumor suppressor RASSF1A preferentially associates with expanded lattice states, revealing differential lattice sensitivity among closely related C1 domain–containing proteins. These short, curved assemblies are enriched at MT intersections and discrete MT segments, revealing spatially heterogeneous lattice states within individual microtubules. At substoichiometric levels, compact lattice–binding proteins behave as both MT compactors and curvature sensors. Changes in osmotic pressure selectively promote dissociation of compact lattice–binding proteins, whereas expanded lattice–binding proteins remain largely unaffected. Using curved filament formation as an in-cellulo readout of compact lattice regions, we identify widespread lattice-state sensitivity across diverse MT-associated and signaling proteins. Finally, we show that these principles extend to neurons, where somatic, but not axonal, tau exhibits sensitivity to lattice expansion despite the expanded lattice architecture of distal axonal microtubules, suggesting additional neuron-specific regulation of lattice accessibility. Together, our findings identify the MT lattice as a dynamic mechanochemical platform whose nanoscale structural states spatially organize protein recruitment and signaling in cells and neurons.

## INTRODUCTION

Microtubules (MTs) are dynamic cytoskeletal polymers that provide mechanical support, organize intracellular architecture, and serve as tracks for motor-driven transport, while also functioning as regulators of cellular force and mechanotransduction through control of Rho signaling (Bershadsky et al., 1996; Birkenfeld et al., 2007; Nalbant et al., 2009; Rafiq et al., 2019). Their functional diversity is regulated by multiple mechanisms, including tubulin isoforms, post-translational modifications (PTMs), and interactions with MT-associated proteins (MAPs), collectively referred to as the “tubulin code”(Akhmanova and Kapitein, 2022; Janke and Magiera, 2020; Roll-Mecak, 2020; Verhey and Gaertig, 2007).

More recently, the structural state of the MT lattice has emerged as an additional regulatory layer. GTP hydrolysis induces angstrom-scale lattice compaction, whereas GTP-like lattice conformations, including those stabilized by the slowly hydrolysable GTP analogue GMPCPP and by MT-stabilizing agents such as taxanes, are generally associated with lattice expansion (Alushin et al., 2014; Peet et al., 2018; Zhang et al., 2018). Structural and in vitro reconstitution studies have further shown that many MAPs preferentially bind either compact or expanded lattice states and can themselves remodel lattice spacing (de Jager et al., 2024; Paquette et al., 2025; Siahaan et al., 2022). For example, tau and doublecortin (DCX) preferentially associate with compact lattice states, whereas proteins such as ensconsin preferentially bind expanded lattices (Shen and Ori-McKenney, 2024; Siahaan et al., 2022). These findings suggest that the MT lattice is not a passive scaffold but a structurally adaptive polymer capable of spatially organizing protein recruitment. Although short curved DCX-positive assemblies have previously been described on bent MTs and attributed to curvature sensing or stabilization (Bechstedt et al., 2014; Ettinger et al., 2016; Paquette et al., 2025), their relationship to MT lattice spacing has not been established, nor has it been determined whether similar structures represent a general feature of compact lattice-binding proteins.

MT lattice spacing is also modulated by physiological and pharmacological perturbations. Taxanes promote lattice expansion, whereas other stabilizing agents, including peloruside A and laulimalide, maintain or induce more compact lattice conformations (Estévez-Gallego et al., 2023; Lucena-Agell et al., 2026; Prota et al., 2014). Likewise, changes in osmotic stress alter MT dynamics and the association of lattice-sensitive MAPs (Molines et al., 2022; Shen and Ori-McKenney, 2024). Together, these observations suggest that the physical state of the cytoplasm and the structural state of the MT lattice are mechanically coupled. However, despite increasing evidence for lattice-dependent protein binding in vitro, how these nanoscale structural states regulate protein organization in cells remains poorly understood because methods to directly probe MT lattice state in situ are lacking.

The MT lattice also functions as a signaling platform by recruiting proteins such as the C1 domain-containing RhoA activator GEF-H1 and the tumor suppressor RASSF1A (Choi et al., 2026). Recent work suggests that GEF-H1 preferentially associates with compact lattice states (Meiring et al., 2025), but whether lattice sensitivity is shared across related C1 domain-containing proteins remains unknown. An additional unresolved question is whether lattice preferences defined largely from structural and in vitro reconstitution studies are maintained in neurons. Recent in situ cryo-electron microscopy revealed that distal axonal MTs in human neurons adopt a stable GMPCPP-like expanded lattice despite being GDP bound, in contrast to compact lattices observed in undifferentiated iPS cells (Zehr et al., 2026). These findings suggest that neuronal MTs possess distinct structural organization and raise the possibility that MT-associated proteins may be regulated differently in this specialized cellular context.

Here, we establish a cellular framework for detecting MT lattice state through the formation of short, curved MT-associated filaments that serve as an in-cellulo readout of compact lattice preference. Using this approach, we provide the first systematic characterization of curved filaments as a cellular signature of compact lattice-binding proteins and show that proteins preferring compact lattices selectively relocalize to these domains following lattice expansion, whereas expanded lattice-binding proteins remain associated with linear MTs and do not accumulate within curved regions. We further demonstrate that this framework distinguishes lattice preferences across diverse structural and signaling proteins, including C1 domain-containing factors, and extend these principles to neurons, where somatic MTs exhibit lattice-dependent protein organization despite the specialized lattice architecture of axons. Together, our findings identify MT lattice spacing as a previously underappreciated mechanochemical regulatory layer that spatially organizes protein recruitment, signaling, and cytoskeletal architecture in cells.

## RESULTS

### Taxol-induced MT lattice expansion promotes curved filaments of tau and doublecortin that are suppressed upon MT lattice compaction

To examine whether MT lattice spacing can be visualized in cells, we treated HeLaM cells with the MT-stabilizing agent Taxol, which has been shown to induce lattice expansion in vitro and in cryo-electron microscopy studies (de Jager et al., 2024; Paquette et al., 2025; Zhang et al., 2018). We then asked whether lattice expansion produces detectable changes in the localization and organization of MT-associated proteins previously shown to preferentially associate with compact MT lattices. To this end, we independently expressed EGFP-tagged tau (human 2N4R) or EGFP–DCX in HeLaM cells (Figure 1), as well as in COS-7 and hTERT RPE1 cells (Supplementary Figure 1A-D). Under control conditions, EGFP–tau displayed the expected filamentous distribution, uniformly decorating MTs (Figure 1A). In contrast, upon Taxol treatment, tau rapidly dissociated from linear MTs and redistributed into numerous short, curved filament-like structures, together with smaller punctate structures (Figure 1B and B’).

**Figure 1.**
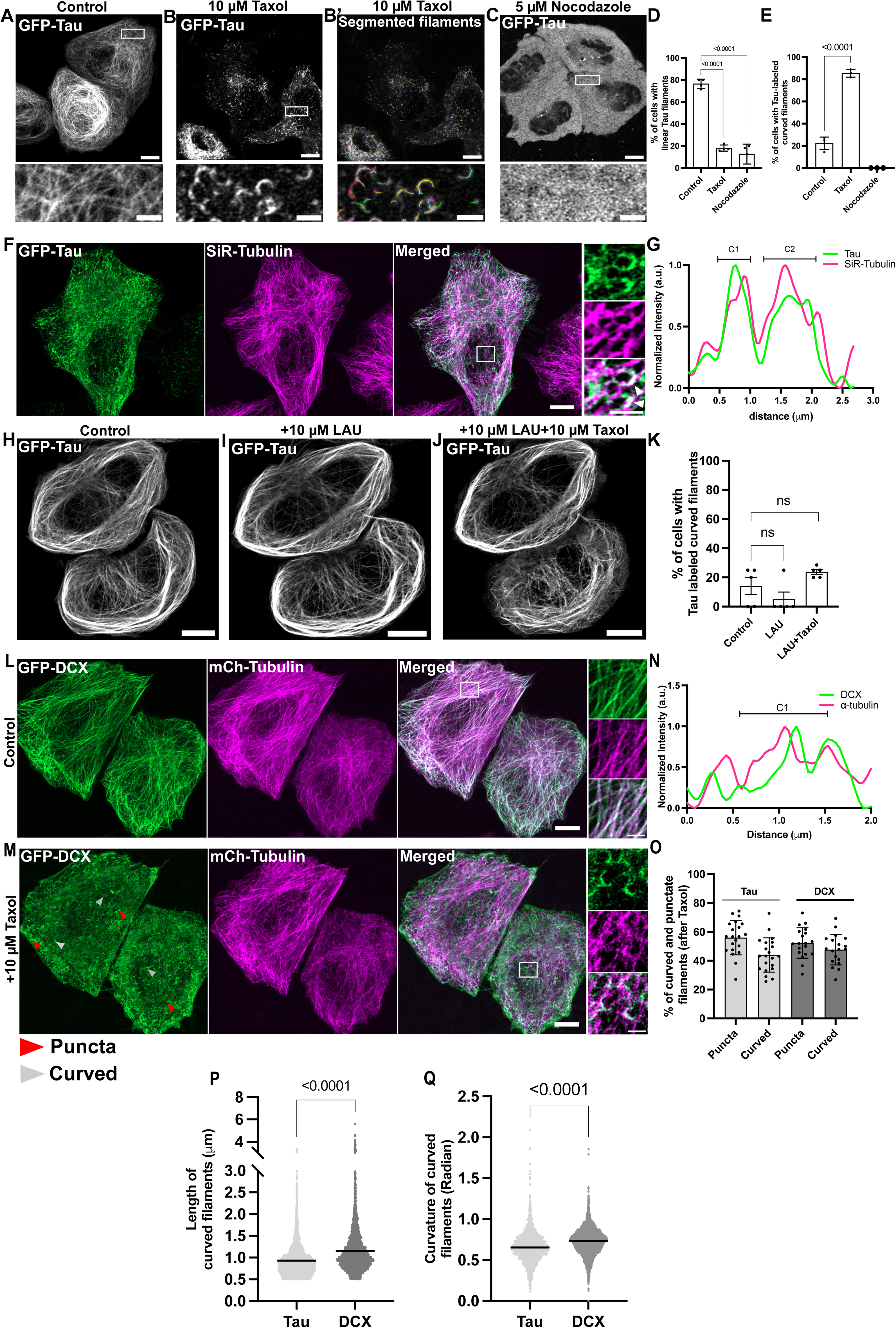
Tau and DCX form short curved MT-associated structures following MT lattice expansion. (A–C) Representative fluorescence images of HeLaM cells expressing EGFP–tau (human 2N4R) under control conditions (A), following treatment with 10 μM Taxol (B), or 5 μM nocodazole (C). Taxol induces redistribution of tau from linear MTs into short curved filament-like and punctate structures. These structures are not observed following nocodazole treatment, indicating that their formation requires an intact MT network. (B’) Automated segmentation of the Taxol-treated cell shown in (B), identifying curved filament-like and punctate tau-positive structures. (D, E) Quantification of cells displaying predominantly linear MT-associated tau filaments (D) or short curved filament-like structures (E) under the indicated treatments (n = 3 independent experiments). ns, not significant; ****, P ff 0.0001 (Welch’s t-test). (F) Representative fluorescence image of a HeLaM cell expressing EGFP–tau (green) and labeled with high-concentration SiR-tubulin (500nM, magenta), showing overlap of tau-positive curved filaments with MTs (see also Supplementary Video 1). Scale bars, 10 μm; inset, 2 μm. (G) Line-scan intensity profile of the representative curved filament shown in (F), demonstrating overlap between EGFP–tau and SiR-tubulin. (H–J) Representative fluorescence images of HeLaM cells expressing EGFP–tau under control conditions (H), following pre-treatment with the MT-compacting agent laulimalide (10 μM; I), or sequential treatment with laulimalide for 30 mins and Taxol (J). Pre-treatment with laulimalide prevents Taxol-induced curved filament formation. (K) Quantification of the percentage of cells displaying tau-positive curved filament-like structures under the indicated treatments (n = 3 independent experiments). ns, not significant; ****, P ff 0.0001 (Welch’s t-test). (L, M) Representative fluorescence images of HeLaM cells co-expressing EGFP–DCX (green) and mCherry–tubulin (magenta) under control (L) and Taxol-treated (M) conditions (see also Supplementary Video 2). Taxol induces curved filament-like and punctate DCX-positive structures that remain associated with MTs. Red arrowheads indicate punctate structures; gray arrowheads indicate curved filaments. Scale bars, 10 μm; inset, 2 μm. (N) Line-scan intensity profile of the representative curved filament shown in (M), demonstrating overlap between EGFP–DCX and mCherry–tubulin. (O–Q) Quantification of the percentage of curved filament-like and punctate structures formed by DCX and tau (O), together with measurements of curved filament length (P) and curvature (Q) (n = 3 independent experiments). ns, not significant (Student’s t-test).

These structures were not observed following MT depolymerization with nocodazole (Figure 1C–E), indicating that their formation requires an intact MT network. Moreover, the curved filament-like structures remained associated with MTs, as revealed by co-labeling with SiR-tubulin (Figure 1F and G) or mCherry–tubulin (Supplementary Figure 1E–H). Notably, high concentrations of SiR-tubulin (>500 nM), which stabilizes MTs through a taxane-related mechanism, similarly induced curved filament formation, whereas lower concentrations (100 nM) did not (Figure 1F; Supplementary Figure 1I-J; Supplementary Videos 1 and 2). Importantly, these curved structures are defined by the selective accumulation of compact lattice–binding proteins rather than by the underlying MT morphology itself, as the entire MT network remains labeled by SiR-tubulin or mCherry–tubulin.

Importantly, similar curved filament-like structures were also observed in cells expressing low levels of tau (based on fluorescence intensity), albeit at lower frequency (Supplementary Figure 1K-L, Figure 1F and G), and became more prominent following Taxol treatment (Figure 1B). These observations suggest that, whereas high tau expression may contribute to local lattice remodeling, lower levels of tau preferentially accumulate within these short, curved MT-associated structures, which may represent localized regions of lattice compaction. To directly test whether the formation of taxol-induced curved filaments depend on MT lattice expansion, we pre-treated cells with laulimalide, a MT-stabilizing agent known to promote or maintain a compact lattice state and to resist Taxol-induced lattice expansion (Estévez-Gallego et al., 2023; Lucena-Agell et al., 2026; Prota et al., 2014). Under these conditions, subsequent Taxol treatment in the presence of laulimalide, failed to induce curved filament formation (Figure 1H-K). Instead, tau remained associated with linear MTs and displayed a uniform decoration pattern.

We next examined whether this behavior extends to other compact lattice–binding proteins. Consistent with tau, expression of EGFP–DCX resulted in the formation of similar curved filament-like and punctate structures upon Taxol treatment (Figure 1L-N), a phenomenon previously reported for DCX (Bechstedt et al., 2014; Ettinger et al., 2016; Paquette et al., 2025). Like tau, co-labeling of EGFP–DCX with mCherry–tubulin confirmed that these structures remained associated with MTs (Figure 1M and N), indicating that they represent MT-bound assemblies rather than detached aggregates.

Quantitative analysis revealed that Taxol treatment induced both curved filament-like and punctate structures in cells expressing either tau or DCX. Curved filaments accounted for 44.01 ± 11.88% and 47.65 ± 10.53% of all structures in tau- and DCX-expressing cells, respectively, whereas punctate structures accounted for 55.99 ± 11.88% and 52.35 ± 10.53% (Figure 1O). Moreover, the curved filaments exhibited defined lengths (DCX: 1.15 ± 0.49 µm; tau: 0.93 ± 0.36 µm) and curvatures (DCX: 0.72 ± 0.16 radian; tau: 0.65 ± 0.18 radian), with tau- and DCX-positive structures displaying highly comparable distributions (Figure 1P and Q). Similar curved filament-like structures were also observed in COS-7 and RPE1 cells expressing either EGFP–tau or EGFP–DCX (Supplementary Figure 1A-D), indicating that lattice expansion–induced relocalization of compact lattice–binding proteins is not restricted to a single cell type. Collectively, these observations suggest that Taxol-induced lattice expansion does not simply alter global MT decoration but instead generates localized MT domains that selectively recruit compact lattice–binding proteins into curved MT-associated assemblies of defined length and curvature.

### Short, curved filaments of DCX and tau localize to segments of individual bent MTs or at MT intersections

To characterize the curved filaments and their spatial distribution relative to the MT architecture, we analyzed both the filaments and the underlying MT network following Taxol-induced stabilization and lattice expansion. Given that the formation of these structures depends on an intact MT polymer, we asked whether curved filaments preferentially localize to specific MT geometries. Consistent with previous reports describing localization to individual bent MTs(Ettinger et al., 2016; Paquette et al., 2025), a subset of curved filaments aligned along segments of single MTs (Figure 2A–C; DCX: 45.44±15.28%; tau: 49.17±16.33%; gray arrowheads). In addition, we frequently detected curved filaments at intersections between overlapping MTs (Figure 2A–C; DCX: 54.56±15.28%; tau: 62.46±15.82%; red arrowheads), indicating that these sites represent prominent regions for their accumulation. Together, these observations suggest that local geometric constraints, including MT bending and filament intersections, may create structural environments that facilitate the emergence or stabilization of compact lattice domains within an otherwise expanded MT network.

**Figure 2.**
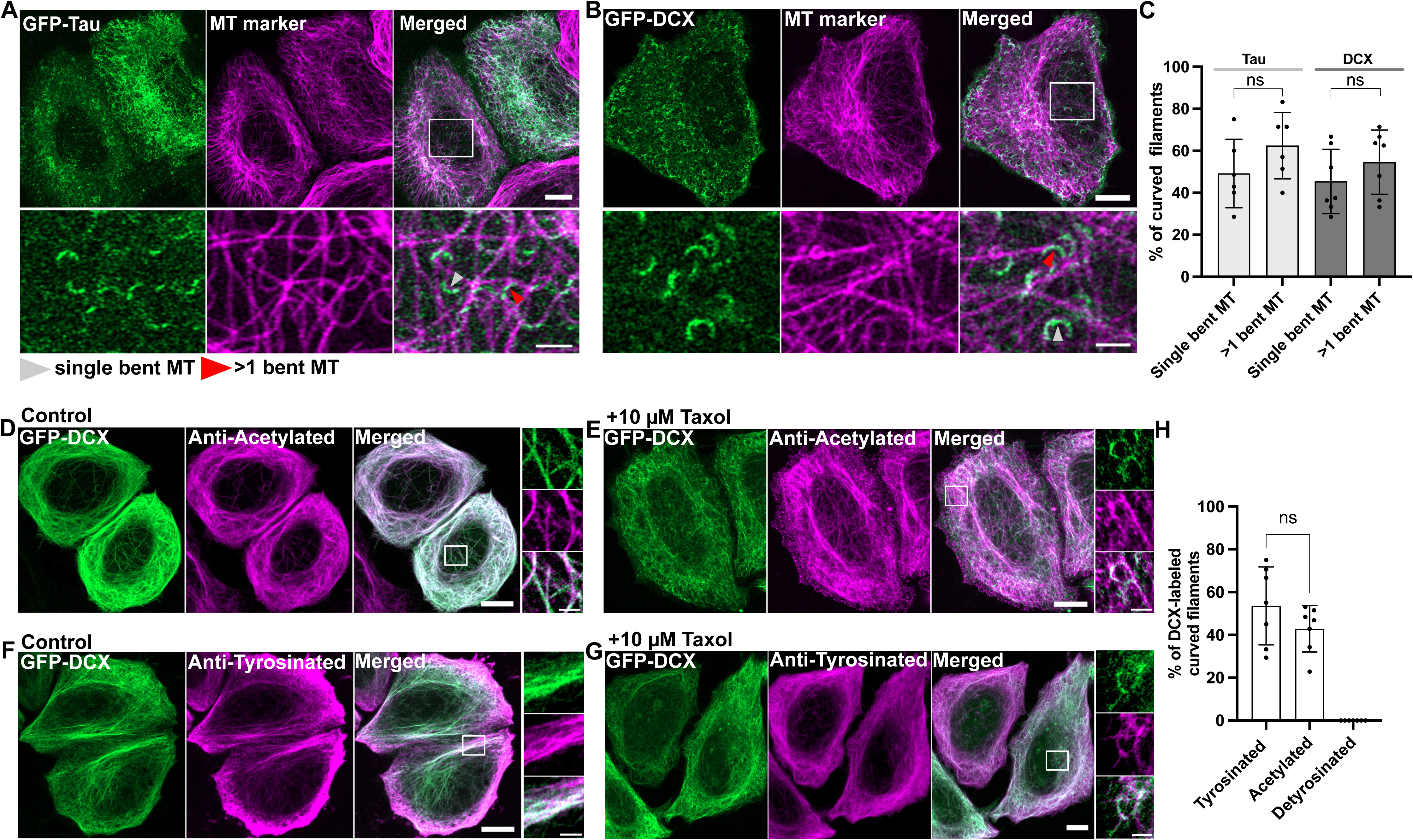
Curved MT-associated structures localize to bent MT segments and MT intersections. (A,B) Representative fluorescence images of HeLaM cells expressing EGFP–tau (A) or EGFP–DCX (B), together with an MT marker (mCherry–tubulin or low concentrations of SiR–tubulin), treated with taxol, showing short curved filaments on bent MTs (gray) or at intersections of more than one MTs (red). (C) Quantification of the proportion of curved filaments localized to bent MTs or MT intersections for tau and DCX (n = 3 independent experiments) ns, not significant (One-way ANOVA). (D–G) Representative fluorescence images of HeLaM cells expressing EGFP–DCX (green) under control (D,F) or taxol-treated (E,G) conditions and stained for acetylated tubulin (D,E, magenta) or tyrosinated tubulin (F,G, magenta). Short curved filaments were associated with acetylated and tyrosinated MTs. Detyrosinated tubulin staining was also performed, but HeLaM cells did not contain detectable detyrosinated MTs (Supplementary Fig. 2A,B). Quantification of the proportion of curved filaments positive for each MT post-translational modification tested (n = 3 independent experiments). ns, not significant (Welch’s t-test). Acetylated, tyrosinated, and detyrosinated tubulin were detected by immunofluorescence using specific antibodies. Scale bars, 10 μm; insets, 2 μm.

To further characterize the molecular properties of curved filament–associated regions, we examined their relationship with established MT post-translational modifications (PTMs), specifically tyrosination, detyrosination, and acetylation. Following Taxol treatment, DCX-positive curved filaments were observed on both acetylated and tyrosinated MTs (Figure 2D–G), indicating that their formation is not restricted to a specific PTM-defined MT population. Notably, HeLaM cells exhibited no detectable detyrosinated MTs, as previously reported (Mohan et al., 2018) and confirmed here (Supplementary Figure 2A). To exclude technical limitations in antibody recognition, we validated detyrosinated MT staining using two independent antibodies in COS-7 cells, both of which robustly detected detyrosinated MTs (Supplementary Figure 2B and C). These observations indicate that detyrosination is not required for curved filament formation, as these structures readily emerged in HeLaM cells despite the absence of detectable detyrosinated MTs. Together, these findings indicate that curved filament formation is not solely determined by the canonical MT PTMs examined here, including acetylation, tyrosination, and detyrosination. Instead, curved filament localization appears to correlate more closely with local structural features of the MT lattice.

Although we cannot exclude contributions from other tubulin modifications, tubulin isotypes, or additional regulatory mechanisms, our findings are consistent with recent work showing that compact lattice-associated recruitment of GEF-H1 occurs independently of these canonical PTMs(Meiring et al., 2025).

### Curved filament formation is specific to compact lattice-binding proteins in the presence of expanders

We next examined how compact lattice–binding proteins behave in a more physiological context, where they coexist with other MAPs, and how the presence of both compact- and expanded lattice–binding proteins influences their organization along MTs. To this end, we co-expressed EGFP–tau and mCherry–ensconsin (MAP7) in HeLaM cells. Expression of the expanded lattice–binding protein ensconsin alone was sufficient to induce multiple distinct tau localization patterns. First, tau-positive filaments frequently did not overlap with ensconsin-labeled MTs, indicating segregation between compact- and expanded lattice–binding proteins (Figure 3A). Second, short curved tau-positive filaments were observed even in the absence of drug treatment, suggesting that expression of an MT lattice expander is sufficient to promote their formation (Figure 3A). Third, in a subset of cells, tau and ensconsin localized to the same MT but occupied alternating, non-overlapping domains (Figure 3B and C), consistent with previous in vitro observations of spatial segregation between these MAPs (Monroy et al., 2018). These results suggest that individual MTs can contain spatially distinct lattice states capable of selectively recruiting compact- or expanded lattice–binding proteins.

**Figure 3.**
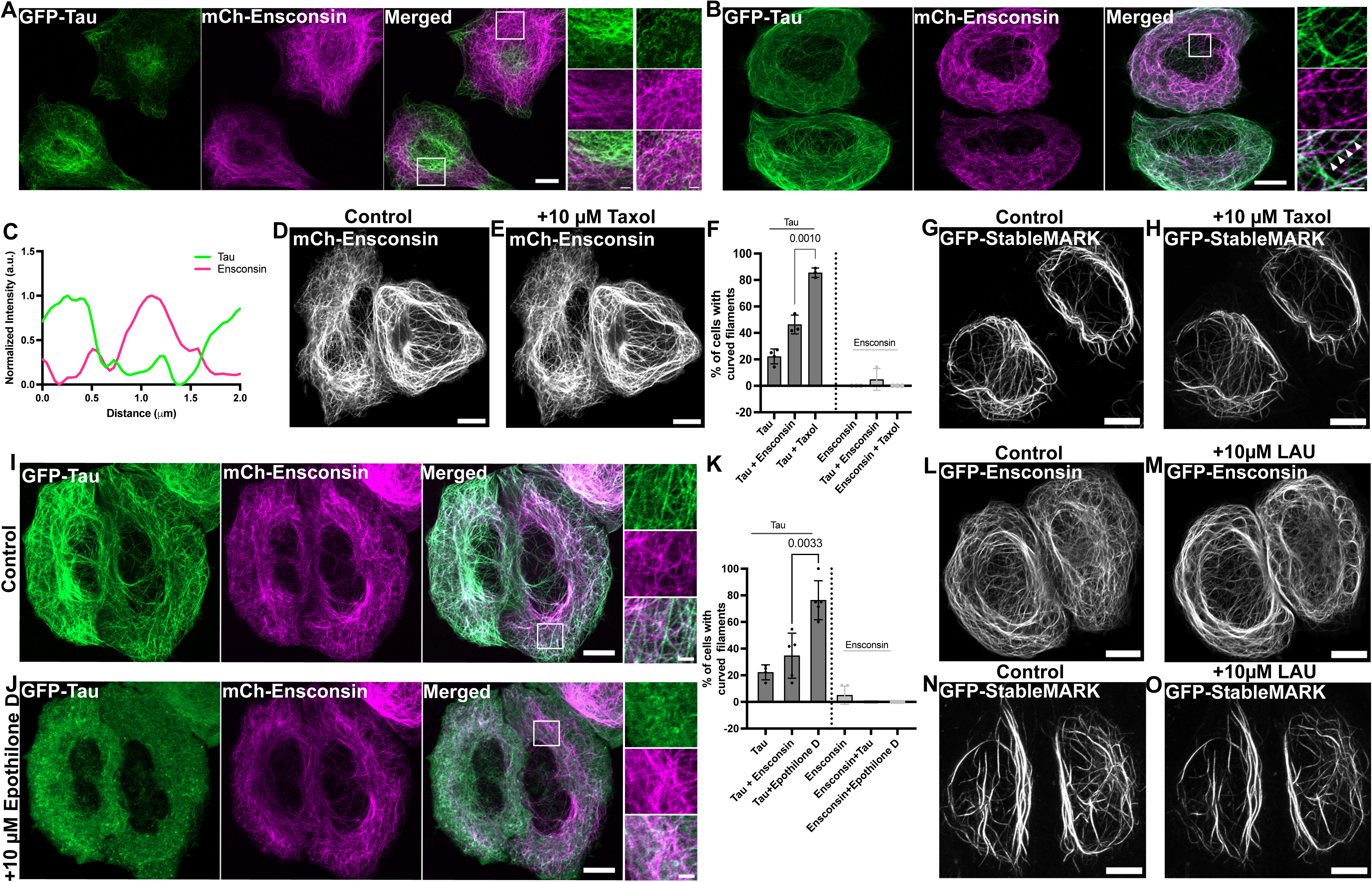
Compact and expanded lattice-binding proteins display distinct responses to MT lattice expansion. (A, B) Representative fluorescence images of HeLaM cells co-expressing EGFP–tau (green) and mCherry–ensconsin (magenta). Distinct populations of tau- and ensconsin- positive MTs are observed on separate filaments (left inset). In addition, the right inset shows spontaneous tau-positive curved filament-like structures in the presence of ensconsin. (B) In some cases, both EGFP-tau (green) and mCherry–ensconsin (magenta) can be observed on the same MT, suggesting the coexistence of compact and expanded lattice states. Scale bars, 10 μm; insets, 2 μm. (C) Line-scan intensity profile of the region indicated by the red arrows in (B), demonstrating alternating localization of EGFP–tau (green) and mCherry–ensconsin (magenta) along the same MT. (D) Representative fluorescence images of HeLaM cells expressing mCherry–ensconsin under control conditions or following Taxol treatment. Unlike tau, ensconsin remains uniformly associated with MTs and does not form curved filament-like structures following Taxol treatment. (E, F) Quantification of the percentage of cells displaying curved filament-like structures under the indicated conditions (n = 3 independent experiments). ns, not significant; ****, P ff 0.0001 (Student’s t-test). (G, H) Representative fluorescence images of HeLaM cells expressing the kinesin-1 rigor construct StableMARK (EGFP–StableMARK) under control conditions (G) or following Taxol treatment (H). Similar to ensconsin, StableMARK remains associated with MTs and does not form curved filament-like structures. (I, J) Representative fluorescence images of HeLaM cells co-expressing EGFP–tau (green) and mCherry–ensconsin (magenta) under control conditions (I) or following treatment with the MT lattice expander epothilone D (J). Epothilone D induces curved filament-like structures of tau but not ensconsin (see also Supplementary Video 3). (K) Quantification of the percentage of cells displaying curved filament-like structures under the indicated treatments (n = 3 independent experiments). ns, not significant; ****, P ff 0.0001 (Welch’s t-test). (L–O) Representative fluorescence images of HeLaM cells expressing EGFP–ensconsin (L,M) or EGFP–StableMARK (N,O) under control conditions or following treatment with the MT-compacting agent laulimalide (10 μM). Neither ensconsin nor StableMARK dissociates from MTs or redistributes into a cytoplasmic pool following laulimalide treatment. Scale bars, 10 μm.

To determine whether curved filament formation is specific to compact lattice–binding proteins, we examined the response of ensconsin and kinesin-1 (StableMARK), a rigor mutant that binds MTs without motility and promotes MT lattice expansion (de Jager et al., 2024; Jansen et al., 2023; Peet et al., 2018). In contrast to tau, neither ensconsin nor StableMARK formed curved filament-like structures following Taxol treatment (Figure 3D–H). Similarly, treatment with the alternative MT-stabilizing agent epothilone D, which also promotes lattice expansion (Lucena-Agell et al., 2026), induced robust formation of tau-positive curved filaments but did not trigger comparable relocalization of ensconsin (Figure 3I–K; Supplementary Video 3) or StableMARK (Supplementary Figure 3A-C). Notably, ensconsin and StableMARK displayed extensive overlap along the same MTs (Supplementary Figure 3D-E), consistent with their shared preference for expanded lattice states. We next asked whether compact lattice stabilization influences the behavior of expanded lattice–binding proteins. In contrast to tau and DCX, treatment with the MT-compacting agent laulimalide did not induce dissociation of either ensconsin or StableMARK from MTs (Figure 3L-O). These observations suggest that expanded lattice–binding proteins remain associated with MTs following laulimalide treatment and are resistant to compaction-induced redistribution. Collectively, these results demonstrate that curved filament formation is a feature specific to compact lattice–binding proteins and establish these structures as a cellular readout of lattice preference. More broadly, the coexistence of alternating tau- and ensconsin-positive domains along individual MTs indicates that structurally distinct lattice states can coexist within the same polymer, providing a potential mechanism for spatial organization of MT-associated proteins in cells.

### Osmotic stress promotes curved filament formation of compact lattice-binding proteins

Changes in osmotic pressure, which may affect cytoplasmic crowding and viscosity, directly regulate MT dynamics by proportionally altering rates of polymerization and depolymerization (Molines et al., 2022). Moreover, osmotic changes has been shown to influence the binding behavior of MT-associated proteins with distinct lattice preferences (Shen and Ori-McKenney, 2024). In particular, hyperosmotic treatments promote the association of expanded lattice–preferring proteins, such as ensconsin and kinesin-1 rigor (StableMARK). We therefore asked whether osmotic modulations in cells could similarly influence the behavior of compact lattice–binding proteins and promote their redistribution into curved filament-like structures.

To test this, HeLaM cells expressing EGFP–tau were treated with 300 mM sucrose to induce a hyperosmotic increase in the cytoplasm. Under these conditions, EGFP–tau was largely redistributed to a diffuse cytoplasmic pool, while the underlying MT network remained intact, as confirmed by SiR-tubulin labeling (Figure 4A and B). Notably, hyperosmotic treatment resulted in elevated background fluorescence and partial cell shrinkage, which often obscured detection of curved tau-positive structures. However, in cells with sufficient signal-to-noise, short, curved filament-like structures were still detectable following sucrose treatment (Figure 4C and D), indicating that these assemblies persist under hyperosmotic conditions. In contrast, treatment with hypotonic medium (0.5x; media diluted 1:1 with water) did not induce additional tau recruitment to MTs, suggesting that reduction of cytoplasmic crowding and/or its viscosity is insufficient to further enhance association of compact lattice–binding proteins. Likewise, hypoosmotic treatment did not induce dissociation of ensconsin, indicating that expanded lattice–binding proteins remain stably associated with MTs across a range of cytoplasmic conditions. Together, these observations indicate that hyperosmotic treatment selectively promotes dissociation of compact lattice–binding proteins (Figure 4A–D), whereas hypoosmotic treatment does not reciprocally enhance their MT association or disrupt expanded lattice–binding proteins (Figure 4E-G).

**Figure 4.**
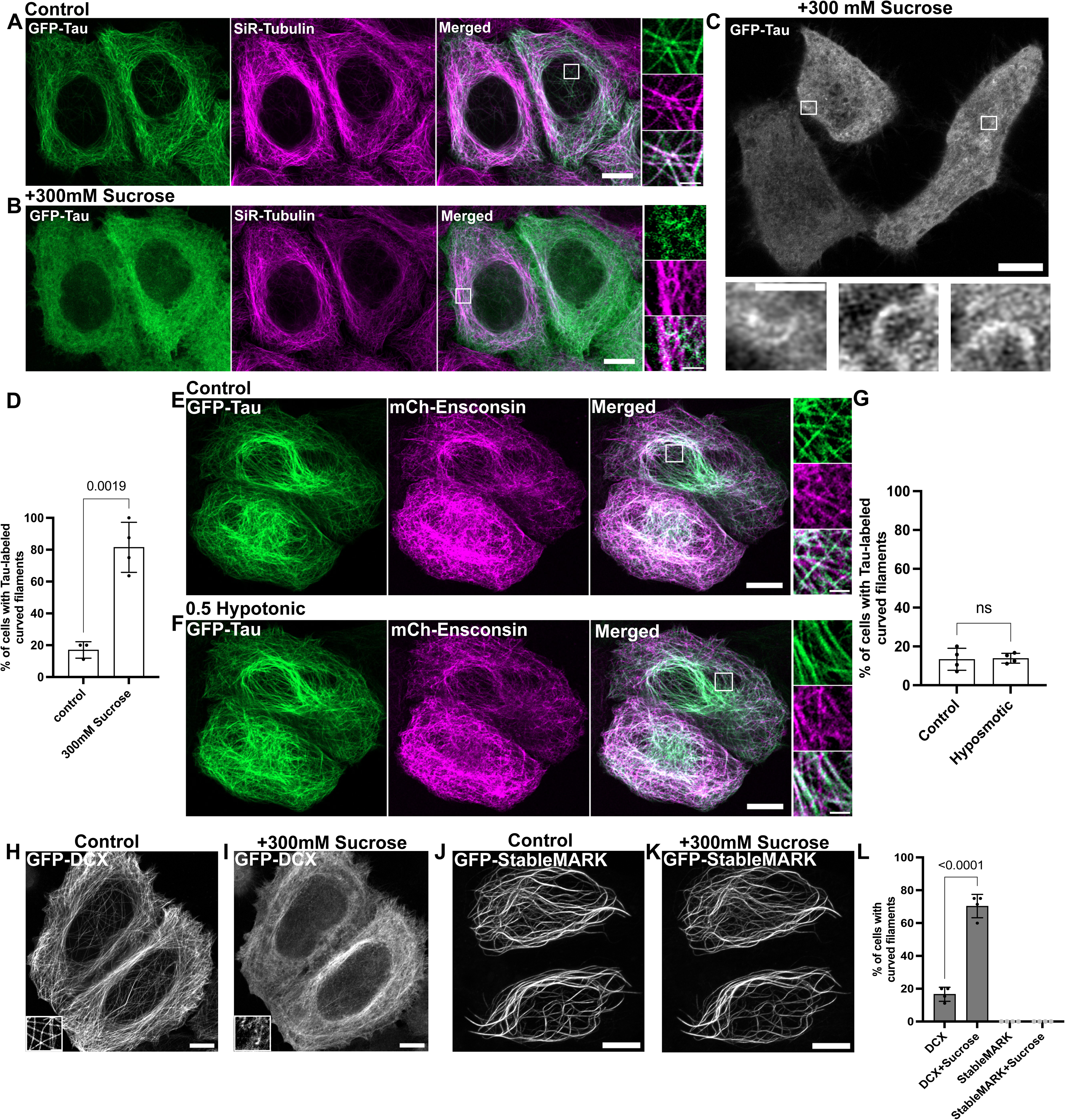
Hyperosmotic stress selectively redistributes compact lattice-binding proteins. ((A–C) Representative fluorescence images of HeLaM cells expressing EGFP–tau (green) and labeled with a low concentration of SiR-tubulin to label MTs (100 nM; magenta) under control conditions (A) or following hyperosmotic treatment with 300 mM sucrose (B, C). Hyperosmotic treatment induces dissociation of EGFP–tau from MTs and its redistribution into curved filament-like structures within 15 min, while the underlying MT network remains intact. Because hyperosmotic treatment causes cell shrinkage, curved filaments are more difficult to visualize; however, cells with a high signal-to-noise ratio of EGFP-tau clearly show their formation (C). Scale bars: (A, B), 10 μm; insets, 2 μm; (C), 5 μm; inset, 1 μm. (D) Quantification of the percentage of cells displaying curved filament-like structures under control and hyperosmotic conditions (n = 3 independent experiments). ns, not significant; ****, P ff 0.0001 (Welch’s t-test). (E, F) Representative fluorescence images of HeLaM cells co-expressing EGFP–tau (green) and mCherry–ensconsin (magenta) under control and hypotonic conditions (1:1 medium). Hypotonic treatment does not induce increased tau association with MTs or dissociation of ensconsin, indicating that reduced cytoplasmic viscosity does not promote MT compaction or release of expanded lattice–binding proteins. Scale bars, 10 μm; insets, 2 μm. (G) Quantification of the percentage of cells displaying curved filament-like structures under hypotonic conditions (n = 3 independent experiments). ns, not significant (Welch’s t-test). (H–K) Representative fluorescence images of HeLaM cells expressing EGFP–DCX (H, I) or EGFP–StableMARK (J, K) following treatment with 300 mM sucrose. Hyperosmotic treatment induces dissociation and curved filament formation of DCX (I) but not StableMARK (K), indicating preferential regulation of compact lattice–binding proteins. (L) Quantification of the percentage of cells displaying curved filament-like structures under the indicated treatments (n = 3 independent experiments). ns, not significant; ****, P ff 0.0001 (Welch’s t-test).

Consistent with tau, EGFP–DCX, another compact lattice–binding protein, similarly dissociated from MTs and formed curved filament-like structures upon sucrose treatment (Figure 4H and I). In contrast, StableMARK, an expanded lattice–preferring protein previously shown to exhibit enhanced MT association under hyperosmotic treatment (Shen and Ori-McKenney, 2024), remained robustly associated with MTs following sucrose treatment and did not display cytoplasmic redistribution or curved filament formation (Figure 4J–L).

### C1 domain–containing signaling proteins display distinct MT lattice preferences

MTs function not only as structural elements but also as signaling platforms that regulate pathways including RhoA, JNK, and stress signaling through direct interactions with signaling proteins. Recent structural studies demonstrated that several C1 domain–containing proteins bind MTs (Choi et al., 2026), while GEF-H1 was recently shown to preferentially associate with compact MT lattices(Meiring et al., 2025). We therefore asked whether our curved filament assay could distinguish lattice preferences of C1 domain–containing signaling proteins directly in cells.

To address this, we examined the C1 domain–containing proteins GEF-H1, RASSF1A, RASSF1D, and PKCζ in HeLaM cells (Figure 5A–F). GEF-H1, RASSF1A, and RASSF1D all exhibited a filamentous MT-like distribution, whereas PKCζ remained largely cytoplasmic (Girik et al., 2024), consistent with previous reports despite the high structural homology among the C1 domains of these proteins (Choi et al., 2026).

**Figure 5.**
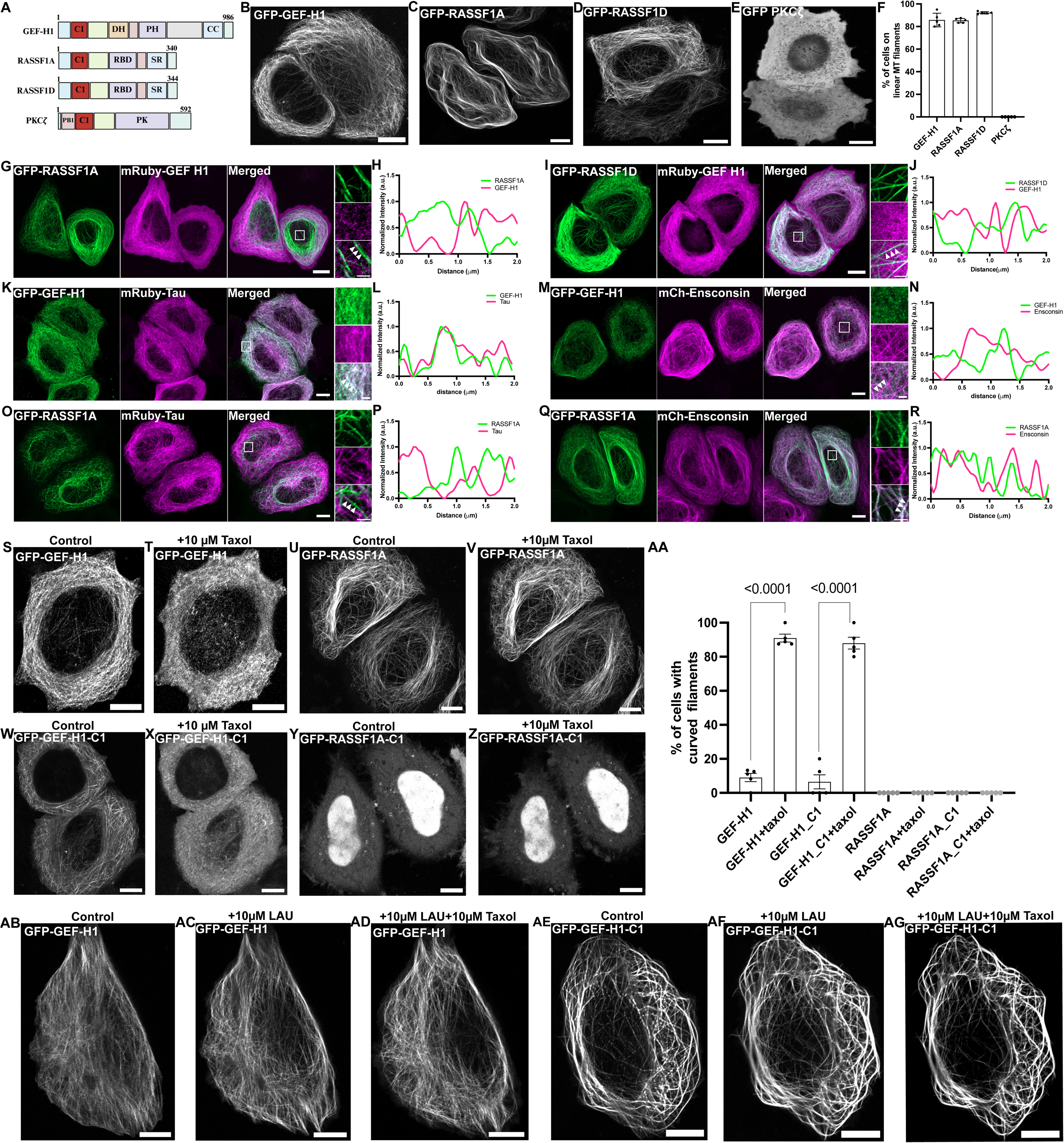
C1 domain-containing signaling proteins exhibit distinct MT lattice preferences. (A) Schematic domain organization of four C1 domain–containing proteins predicted and/or reported to associate with MTs: GEF-H1, RASSF1A, RASSF1D, and PKCζ. The C1 domain is highlighted in red. (B–E) Representative fluorescence images of HeLaM cells expressing EGFP–GEF-H1 (B), EGFP–RASSF1A (C), EGFP–RASSF1D (D), or EGFP–PKCζ (E). EGFP–GEF-H1, EGFP–RASSF1A, and EGFP–RASSF1D display filamentous MT-like localization, whereas PKCζ remains largely cytoplasmic despite the high structural homology of its C1 domain with those of GEF-H1 and RASSF1. Scale bars, 10 μm. (F) Quantification of the percentage of cells displaying filamentous localization for the indicated C1 domain–containing proteins (n = 3 independent experiments). (G–J) Representative fluorescence images of HeLaM cells co-expressing EGFP–RASSF1A (green; G) or EGFP–RASSF1D (green; I) together with mRuby–GEF-H1 (magenta). RASSF1 isoforms and GEF-H1 show largely non-overlapping filamentous localization, with increased diffuse cytoplasmic localization of GEF-H1 in the presence of RASSF1 isoforms. (H, J) Line-scan intensity profiles of the regions indicated in (G) and (I). (K–N) Representative fluorescence images of HeLaM cells co-expressing EGFP–GEF-H1 (green; K) or EGFP–RASSF1A (green; O) together with mRuby–tau (magenta). EGFP–GEF-H1 extensively overlaps with tau-positive MTs, whereas EGFP–RASSF1A displays distinct filamentous localization with minimal overlap with tau. (L, P) Line-scan intensity profiles of the indicated regions. (M and Q) Representative fluorescence images of HeLaM cells co-expressing EGFP–GEF-H1 (green; M) or EGFP–RASSF1A (green; Q) together with mCherry–ensconsin (magenta). EGFP–GEF-H1 becomes largely cytoplasmic in the presence of the expanded lattice–binding protein ensconsin (See also Supplementary Video 4), whereas EGFP–RASSF1A shows extensive overlap with ensconsin-positive MTs. (N, R) Line-scan intensity profiles of the indicated regions. Scale bars, 10 μm; insets, 2 μm. (S, T, U, V) Representative fluorescence images of HeLaM cells expressing full-length EGFP–GEF-H1 (S, T) or EGFP–RASSF1A (U, V) under control conditions (S, U) or following Taxol treatment (T, V). Unlike GEF-H1, which becomes cytoplasmic and relocalize to short curved filaments, EGFP–RASSF1A remains associated with MTs following Taxol treatment, indicating a lack of preference for compact MT lattice states. (W, X) Representative fluorescence images of HeLaM cells expressing the isolated EGFP–GEF-H1 C1 domain (aa 28–100; W, X) under control conditions (W) or following Taxol treatment (X, See also Supplementary Video 5). Both full-length GEF-H1 and the isolated C1 domain dissociate from MTs and relocalize into curved filament-like structures following Taxol treatment. (Y, Z) Representative fluorescence images of HeLaM cells expressing the isolated EGFP–RASSF1A C1(aa 1- 110) domain under control conditions (Y) or following Taxol treatment (Z). Unlike the GEF-H1 C1 domain, the isolated RASSF1A C1 domain displays predominantly nuclear- like localization rather than MT association and shows no detectable redistribution following Taxol treatment. Scale bars, 10 μm. (AA) Quantification of the percentage of cells displaying curved filament-like structures under the indicated conditions shown in (S–Z) (n = 3 independent experiments). ns, not significant; ****, P ff 0.0001 (Welch’s t- test). (AB–AG) Representative fluorescence images of HeLaM cells expressing full- length EGFP–GEF-H1 (AB–AD) or the isolated EGFP–GEF-H1 C1 domain (AE–AG) following treatment with laulimalide alone (AC, AF) or sequential treatment with laulimalide and Taxol (AD, AG). Laulimalide prevents Taxol-induced curved filament formation in both full-length GEF-H1 and the isolated GEF-H1 C1 domain, indicating that GEF-H1 preferentially associates with compact MT lattice states through its C1 domain. Scale bars, 10 μm.

RASSF1A is the canonical tumor suppressor isoform, whereas RASSF1D is predominantly expressed in cardiac tissue (Dubois et al., 2019; van der Weyden and Adams, 2007). Although the two isoforms differ only within the C1 domain, with RASSF1D containing an additional RSALD sequence that forms a longer predicted loop (Supplementary Figure 4A and B), both C1 domains remain highly homologous.

Consistent with previous reports, EGFP–GEF-H1 strongly decorated MTs (Supplementary Figure 4C), whereas EGFP–PKCζ showed no detectable MT association (Figure 5E and F). Interestingly, co-expression of either EGFP–RASSF1A or EGFP– RASSF1D with mRuby–GEF-H1 redistributed GEF-H1 into a diffuse cytoplasmic pool (Figure 5G–J), suggesting that stable association of GEF-H1 and RASSF1 isoforms with the same MT lattice is incompatible. Consistent with its reported compact lattice preference, EGFP–GEF-H1 extensively co-localized with mRuby–tau (Figure 5K and L). In contrast, co-expression with the expanded lattice–binding protein mCherry–ensconsin redistributed GEF-H1 into a diffuse cytoplasmic pool and, in a subset of cells, into short curved filament-like structures (Figure 5M and N; Supplementary Video 4), indicating preferential association of GEF-H1 with compact lattice regions.

We next asked whether this behavior differed from that of other C1 domain–containing proteins. Both RASSF1A and RASSF1D displayed localization patterns distinct from GEF-H1. Rather than co-localizing with tau-positive MTs, both isoforms preferentially associated with ensconsin-labeled MTs (Figure 5O–R; Supplementary Figure 4D-F).

Unlike GEF-H1, which dissociated from MTs and formed curved filament-like structures following Taxol treatment (Figure 5S and T), RASSF1A and RASSF1D remained stably associated with MTs and showed no detectable curved filament formation (Figure 5U and V; Supplementary Figure 4D and E). These observations indicate that, despite containing highly homologous C1 domains, GEF-H1 preferentially associates with compact lattice states, whereas RASSF1 isoforms preferentially associate with expanded lattice states.

To determine whether these differences are encoded within their C1 domains, we examined isolated C1-domain constructs. Taxol induced robust curved filament formation in both full-length GEF-H1 and its isolated C1 domain (aa 28–100) (Figure 5W and X; Supplementary Video 5), demonstrating that the GEF-H1 C1 domain is sufficient to confer lattice sensitivity. In contrast, although deletion of the C1 domain abolishes MT association of RASSF1A (Choi et al., 2026), the isolated C1 domains of both RASSF1A (aa 1-110) and RASSF1D (aa 1-114) failed to associate with MTs and instead displayed predominantly cytoplasmic or nuclear localization (Figure 5Y and Z; Supplementary Figure 4G and H). Notably, the four additional amino acids (LSAD) present within the RASSF1D C1 domain did not alter its localization behavior relative to RASSF1A. Importantly, curved filament formation by both full-length GEF-H1 and its isolated C1 domain was completely suppressed by pre-treatment with the MT- compacting agent laulimalide (Figure 5AB–AG), consistent with our observations for tau (Figure 1).

Finally, we examined PKCζ, another C1 domain–containing signaling protein previously reported not to associate with MTs (Choi et al., 2026; Girik et al., 2024). Consistent with these reports, EGFP–PKCζ remained largely cytoplasmic and showed no detectable MT association or lattice-sensitive redistribution (Supplementary Figure 4I). Collectively, these findings demonstrate that lattice sensitivity is not a universal property of C1 domains. Instead, closely related C1 domain–containing signaling proteins exhibit distinct MT lattice preferences, with the GEF-H1 C1 domain being sufficient to confer compact lattice recognition, whereas RASSF1 requires additional determinants for MT association and preferentially associates with expanded lattice states.

### Differential lattice sensitivity of MT-associated proteins revealed by curved filament formation

Since our approach identified distinct lattice-dependent behaviors for compact and expanded lattice–binding proteins, we next sought to determine whether additional MAPs and MT-associated proteins also display selective sensitivity to MT lattice spacing. We therefore expanded our analysis to a broader set of structural and regulatory MAPs using curved filament formation and redistribution behavior as an in-cellulo readout of lattice preference.

Interestingly, although many proteins exhibited clear preferences for either compact or expanded lattice states, several classical structural MAPs appeared largely insensitive to lattice perturbation. For example, MAP4-mApple remained associated with MTs following Taxol treatment and did not form curved filament-like structures (Figure 6A and B). Consistent with these observations, MAP4 has previously been shown to be largely insensitive to lattice spacing in vitro (Siahaan et al., 2022). Similarly, EGFP– MAP1B neither dissociated from MTs nor redistributed into curved filament-like structures following Taxol treatment (Figure 6C and D) and overlapped extensively with both the compact lattice–binding protein tau (Figure 6E and F) and the expanded lattice marker ensconsin (Figure 6G and H). Together, these findings suggest that not all MAPs strongly discriminate between lattice conformations and that some structural MAPs tolerate a broader range of lattice geometries.

**Figure 6.**
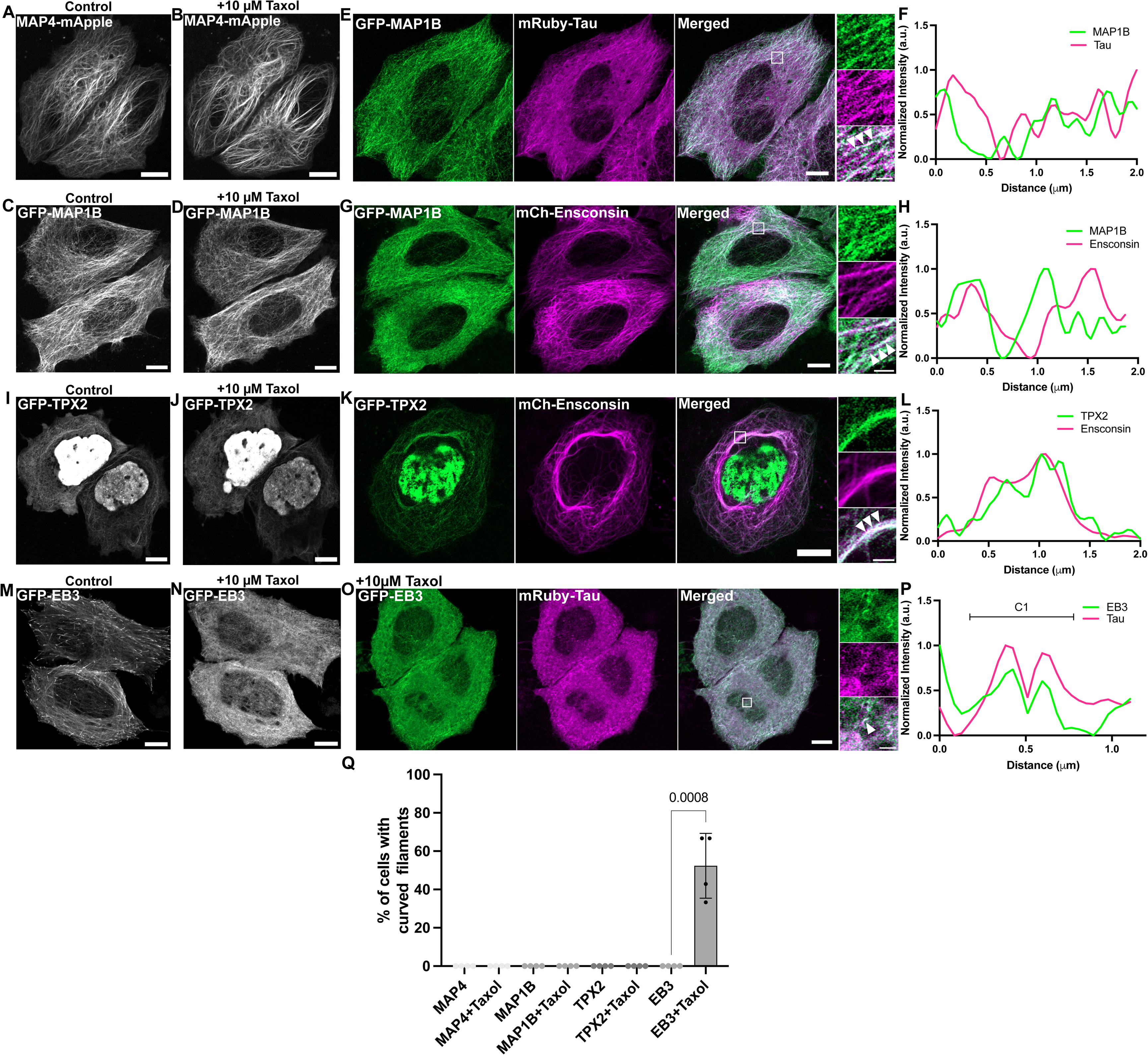
MT-associated proteins exhibit distinct lattice-dependent localization behaviors. (A–D) Representative fluorescence images of HeLaM cells expressing MAP4-mApple (A, B) or EGFP–MAP1B (C, D) under control conditions (A, C) or following Taxol treatment (B, D). Neither MAP4 nor MAP1B shows noticeable changes in MT association or redistribution following Taxol treatment. (E–H) Representative fluorescence images of HeLaM cells co-expressing EGFP–MAP1B (green) together with mRuby–tau (magenta; E, F) or mCherry–ensconsin (magenta; G, H). MAP1B overlaps with both tau- and ensconsin-labeled MTs, suggesting a lack of preference for either compact or expanded MT lattice states. (F, H) Line-scan intensity profiles of the indicated regions. Scale bars, 10 μm; insets, 2 μm. (I–L) Representative fluorescence images of HeLaM cells expressing EGFP–TPX2 under control conditions (I) or following Taxol treatment (J). EGFP–TPX2 localizes to both MTs and the nucleus and remains associated with MTs following Taxol treatment. Co-expression of EGFP–TPX2 (green) with mCherry– ensconsin (magenta) shows extensive overlap on MTs (K). (L) Line-scan intensity profile of the indicated region. (M–O) Representative fluorescence images of HeLaM cells expressing EGFP–EB3 under control conditions (M) or following Taxol treatment (N, O). EGFP–EB3 displays a characteristic comet-like distribution under control conditions but becomes largely cytoplasmic and relocalizes into curved filament-like structures following Taxol treatment (green; O). In Taxol-treated cells, EB3-positive curved structures overlap with mRuby–tau curved filaments (magenta; O). Scale bars, 10 μm; insets, 2 μm. (P) Line-scan intensity profile of the representative curved filament shown in (O), demonstrating overlap between EGFP–EB3 and mRuby–tau. (Q) Quantification of the percentage of cells displaying curved filament-like structures under the indicated conditions (n = 3 independent experiments). ns, not significant; ****, P ff 0.0001 (Student’s t-test).

In contrast, TPX2, a mitotic spindle assembly factor previously proposed to associate preferentially with expanded or stabilized MT lattices (de Jager et al., 2024; Liang et al., 2025; Zhang et al., 2017), remained robustly associated with MTs following Taxol treatment without detectable curved filament formation or cytoplasmic redistribution and strongly overlapped with ensconsin-decorated MTs (Figure 6I–L). These observations further support the idea that expanded lattice–binding proteins remain stably associated with Taxol-expanded MTs.

We next examined EB3, a canonical plus-end tracking protein (+TIP) known to preferentially recognize the GTP lattice at growing MT ends (de Jager et al., 2024; Maurer et al., 2012; Zhang et al., 2018). Consistent with previous studies, EGFP–EB3 localized prominently to growing MT plus ends under control conditions (Supplementary Figure 5). Surprisingly, however, EB3 became largely cytoplasmic following Taxol or Epothilone D treatments and formed curved filament-like assemblies (Figure 6M–P, Supplementary Figure 5C and D), despite them promoting lattice expansion. This observation is consistent with previous work demonstrating that EB proteins recognize a specific nucleotide-dependent lattice geometry associated with the GTP cap rather than an expanded lattice state per se, and that Taxol-expanded lattices are structurally distinct from native EB3-binding conformations (Zhang et al., 2018). Thus, lattice expansion alone appears insufficient to support EB3 association in the absence of the proper GTP-cap architecture.

### MT lattice–dependent protein organization extends to neurons

Because many of the proteins examined in this study are highly enriched in neurons, we next asked whether lattice-dependent protein behavior could also be observed in a neuronal context. To address this, we expressed EGFP–tau in human cortical-like i^3^Neurons generated through Neurogenin-2 induction and analyzed them at days 18–22, a stage at which these neurons form mature neurites and functional synaptic contacts (Fujise et al., 2025). Similar to HeLaM cells, curved filament-like structures became more pronounced following treatment with high concentrations of SiR-tubulin (>500nM; Figure 7C and D), but not low concentrations (50-100nM; Figure 7A and B), suggesting that partial MT stabilization or lattice expansion promotes the formation of compact lattice-associated domains in neurons.

**Figure 7.**
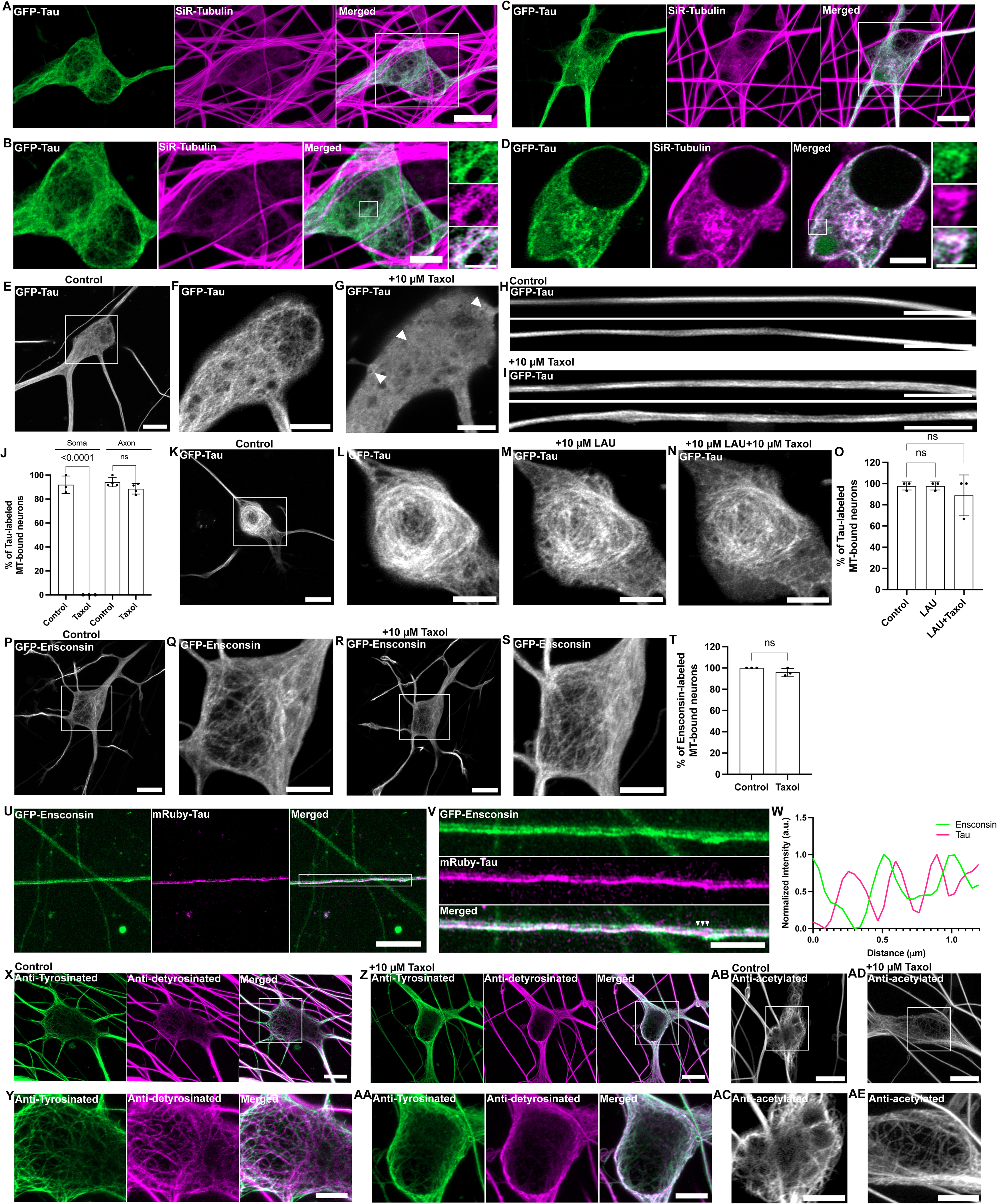
Lattice-dependent protein organization is conserved in the somatic region of neurons. (A–D) Representative fluorescence images of human i^3^Neurons (days 18–22) expressing EGFP–tau (green) and labeled with low concentrations of SiR-tubulin (100 nM; magenta) (A, B; soma) or high concentrations of SiR-tubulin (500 nM; C, D; soma). High concentrations of SiR-tubulin induce spontaneous curved filament-like structures, similar to those observed in HeLaM cells. Scale bars: (A, C), 10 μm; (B, D), 5 μm; insets, 2 μm. (E–G) Representative fluorescence images of i^3^Neurons expressing EGFP–tau under control conditions (E, F; soma) or following Taxol treatment (G). Taxol induces redistribution of tau into a diffuse cytoplasmic pool together with curved filament-like structures within the somatic region. Curved filaments are partially masked by the increased cytoplasmic background. Scale bars: (E), 10 μm; (F, G), 5 μm. (H,I) Representative fluorescence images of two axonal regions from i^3^Neurons expressing EGFP–tau under control conditions (H) or following Taxol treatment (I). In contrast to the soma, tau remains predominantly associated with axonal MTs following Taxol treatment. Scale bars, 5 μm. (J) Quantification of the percentage of neurons displaying MT-associated tau filaments in the soma and axon under control and Taxol-treated conditions (n = 3 independent experiments). ns, not significant; ****, P ff 0.0001 (Student’s t-test). (K–N) Representative fluorescence images of i^3^Neurons expressing EGFP–tau under control conditions (K, L; soma), following pre-treatment with the MT- compacting agent laulimalide (M), or sequential treatment with laulimalide and Taxol (N). Pre-treatment with laulimalide preserves tau association with MTs and prevents the Taxol-induced dissociation observed with Taxol treatment alone. Scale bars: (K), 10 μm; (L–N), 5 μm. (O) Quantification of the percentage of neurons displaying MT-associated tau filaments in the soma under the indicated treatments (n = 3 independent experiments). ns, not significant (Student’s t-test). (P–S) Representative fluorescence images of i^3^Neurons expressing EGFP–ensconsin under control conditions (P, Q; soma), following Taxol treatment (R, S; soma). EGFP–ensconsin remains associated with MTs under all conditions. Scale bars: (P, R), 10 μm; (Q, S), 5 μm. (T) Quantification of the percentage of neurons displaying MT-associated ensconsin filaments in the soma under the indicated treatments (n = 3 independent experiments). ns, not significant (Student’s t-test). (U, V) Representative fluorescence images of i^3^Neurons co- expressing EGFP–Ensconsin (green) and mRuby-Tau(magenta). High-magnification images (V) reveal distinct alternating regions enriched for tau or ensconsin along axonal MTs. Scale bars: (U), 10 μm; (V), 5 μm. (W) Line-scan intensity profile of the region indicated by the white box in (U). (X, Y–AA) Representative fluorescence images of i^3^Neurons immunolabeled for tyrosinated and detyrosinated tubulin under control conditions (X, Y; soma) or following Taxol treatment (Z, AA; soma). Tyrosinated and detyrosinated MTs exhibit similar distributions under both conditions. (AB–AE) Representative fluorescence images of i^3^Neurons immunolabeled for acetylated tubulin under control conditions (AB, AC; soma) or following Taxol treatment (AD, AE; soma). Acetylated MTs likewise show no obvious changes in distribution following Taxol treatment. Acetylated, tyrosinated, and detyrosinated tubulin were detected by immunofluorescence using specific antibodies. Scale bars: (X, Z, AB, AD), 10 μm; (Y, AA, AC, AE), 5 μm.

Treatment with Taxol induced a pronounced redistribution of tau from somatic MTs into a diffuse cytoplasmic pool together with curved filament-like structures (Figure 7E–G), closely resembling our observations in non-neuronal cells. Interestingly, tau association with axonal MTs, identified based on neurite morphology and length, was comparatively resistant to Taxol (Figure 7H-J). This observation is notable in light of a recent structural study demonstrating that axonal MTs in i^3^Neurons predominantly contain GDP-tubulin while adopting a stable GMPCPP-like expanded lattice conformation (Zehr et al., 2026). Importantly, Taxol-induced tau dissociation within the somatic compartment was completely prevented by pre-treatment with the MT-compacting agent laulimalide (Figure 7K–O), indicating that neuronal tau localization, like that in non-neuronal cells, is regulated by MT lattice state. Together, these findings suggest that additional regulatory mechanisms, potentially involving neuronal MAP composition, MT bundling, or axon-specific lattice organization, stabilize tau association within axonal MT arrays despite global lattice expansion. In parallel, EGFP–ensconsin, which preferentially associates with expanded lattice states, remained robustly associated with somatic MTs following Taxol treatment, and did not undergo detectable cytoplasmic redistribution, unlike tau (Figure 7P–T).

To determine whether axonal MTs can simultaneously recruit compact- and expanded lattice-binding proteins, we co-expressed EGFP–tau and mCherry–ensconsin in i^3^Neurons (Figure 7U–W). Both proteins localized to discrete alternating regions along axons, with limited overlap between tau- and ensconsin-positive domains, suggesting preferential association with non-overlapping MT segments within the axonal bundle. Given that axonal MTs are organized into tightly packed bundles, these observations suggest that heterogeneous lattice organization exists within axons. Whether this reflects differences between neighboring microtubules within a bundle or along distinct segments of individual microtubules cannot be resolved at the current imaging resolution.

Finally, we asked whether acute manipulation of MT lattice spacing alters canonical tubulin post-translational modifications (PTMs) in neurons. Co-staining of βIII-tubulin with tyrosinated tubulin confirmed that overall neuronal MT organization remained preserved following Taxol treatment (Supplementary Figure 6A and B). Analysis of tyrosinated, detyrosinated, and acetylated tubulin revealed no major changes in their abundance or distribution following acute Taxol treatment (Figure 7X, Y and AB–AC). Specifically, tyrosinated, detyrosinated, and acetylated MT populations remained readily detectable in both control and Taxol-treated neurons, with no overt redistribution in either the soma (Figure 7Z, AA, AD and AE) or axons (Supplementary Figure 6C–F). Although these observations do not exclude longer-term coupling between MT lattice state and tubulin PTMs, they indicate that acute modulation of MT lattice spacing (30–60 min) can occur independently of major changes in canonical tubulin modifications. Together, these findings extend lattice-state–dependent protein organization to neurons and support the idea that MT structural plasticity contributes to the spatial regulation of neuronal MAPs and signaling proteins within specialized neuronal compartments.

## DISCUSSION

Our findings identify the MT lattice as a dynamic mechanochemical platform whose local structural state spatially regulates the recruitment of MT-associated and signaling proteins in cells. While angstrom-scale differences in MT lattice spacing have been extensively characterized in vitro, whether such structural states could be functionally resolved in cells has remained unclear. Here, we establish short, curved MT-associated filaments as the first in-cellulo readout of compact MT lattice regions and show that proteins previously associated with compact lattices, including tau, DCX, and GEF-H1, selectively relocalize to these domains upon MT lattice expansion. Conversely, maintenance of compact lattice states through laulimalide suppresses this response, supporting the idea that local MT architecture directly regulates protein organization in cells. Although short curved DCX-positive assemblies have previously been described on bent MTs (Bechstedt et al., 2014; Ettinger et al., 2016; Paquette et al., 2025), our work demonstrates that these structures represent a broader feature of compact lattice-binding proteins and can be exploited as a cellular readout of MT lattice preference.

A key observation from our study is that Taxol-induced lattice expansion does not generate a uniformly expanded MT network. Instead, expanded lattices coexist with discrete curved regions enriched in compact lattice-binding proteins, frequently localizing along individual MT segments or at MT intersections. These observations suggest that MTs contain spatially heterogeneous mechanochemical domains rather than existing as structurally uniform polymers. The enrichment of compact lattice-binding proteins at curved MT regions further suggests a close relationship between MT curvature and lattice architecture, although the structural basis of this coupling remains unknown. Collectively, our findings suggest that Taxol-induced lattice expansion does not uniformly remodel the MT network but instead generates localized compact lattice regions along individual MT segments and at MT intersections that selectively recruit compact lattice-binding proteins despite a globally expanded MT state.

Our PTM analyses further indicate that these compact lattice regions cannot be explained solely by canonical tubulin modifications. Previous work has suggested that lattice expansion is associated with increased tyrosination and that tubulin PTMs correlate with distinct lattice conformations (Hotta et al., 2026; Yue et al., 2026). In contrast, curved filaments readily formed in HeLaM cells, which lack detectable detyrosinated MTs, and were associated with both tyrosinated and acetylated MT populations. These observations suggest that canonical PTMs are not sufficient to define the compact lattice regions recognized by tau, DCX, or GEF-H1. Rather, lattice spacing may represent a structural regulatory layer that operates independently of, or in parallel with, tubulin PTMs.

An important question raised by our findings concerns the ultrastructural organization of these curved MT-associated domains. One intriguing possibility is that MT bending generates asymmetric lattice conformations across the wall of an individual MT. Under this model, the convex surface of a curved MT would experience tensile strain and adopt a relatively expanded lattice geometry, whereas the concave surface would undergo compressive strain and locally assume a compact lattice-like conformation (Figure 8). Such local asymmetry could allow compact lattice-binding proteins, such as tau and DCX, to preferentially accumulate on the concave surface, while expanded lattice-binding proteins, including ensconsin, may still associate with the convex surface. Such structural asymmetry could explain how compact lattice-binding proteins, including tau, DCX, and GEF-H1, become enriched within curved MT regions despite an overall expanded lattice state and regardless of the global PTM identity of the MT. Resolving this possibility will require higher-resolution structural approaches, such as cryo-electron tomography or correlative cryo-light and electron microscopy.

**Figure 8.**
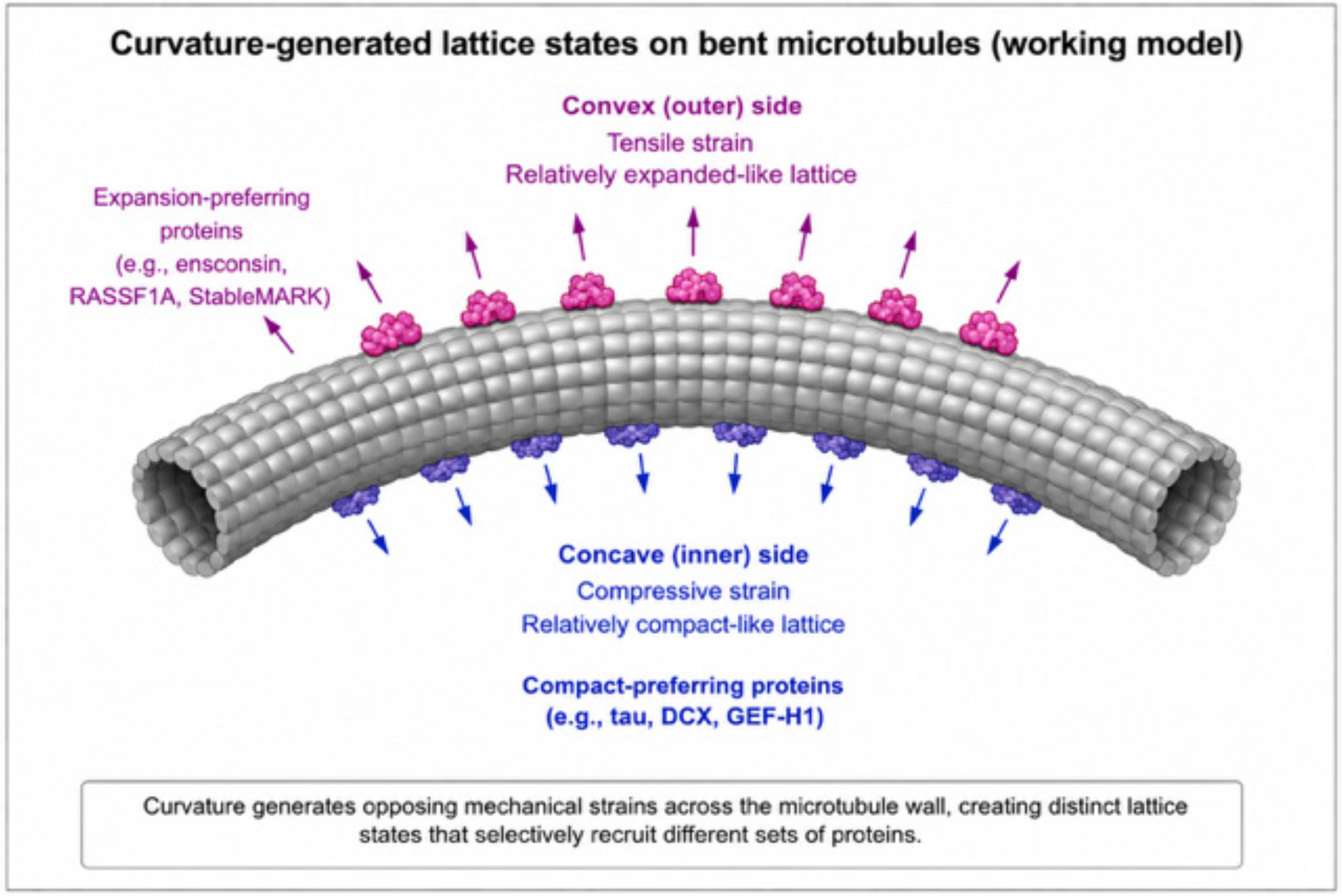
Proposed model for curvature-generated lattice asymmetry in bent microtubules. Schematic illustrating a working model in which microtubule bending generates opposing mechanical strains across the microtubule wall. Under this model, the convex surface experiences tensile strain and adopts a relatively expanded lattice-like conformation that favors recruitment of expanded lattice–binding proteins, whereas the concave surface experiences compressive strain and adopts a relatively compact lattice-like conformation that favors recruitment of compact lattice–binding proteins. This model provides a potential explanation for the enrichment of compact lattice– binding proteins within curved microtubule regions despite an overall expanded lattice state. The illustration was generated with the assistance of artificial intelligence and subsequently edited and curated by the authors.

Our data further suggest that compact- and expanded lattice-binding proteins can coexist on distinct regions of the same MT. Tau and ensconsin frequently segregated onto separate MT populations but could also occupy alternating, non-overlapping domains along individual filaments, indicating that single MTs can contain spatially distinct lattice states. Importantly, lattice expansion selectively displaced compact- binding proteins while preserving expanded lattice-binding proteins such as ensconsin and kinesin-1 rigor (StableMARK), suggesting that different MAPs may stabilize or reinforce specific lattice conformations. Together, these findings support a model in which individual MTs behave as structurally adaptive polymers capable of spatially partitioning protein recruitment through local lattice organization. Changes in osmotic stress further modulated these lattice-dependent behaviors. Hyperosmotic conditions selectively promoted dissociation of compact lattice-binding proteins and enhanced curved filament formation, whereas expanded lattice-binding proteins remained associated with MTs. Although hyperosmotic treatment is known to alter MT polymerization dynamics(Molines et al., 2022), our data suggest that it also remodels the accessibility of distinct lattice states. Whether these effects arise directly through lattice remodeling or indirectly through altered MT dynamics remains to be determined.

An important implication of our work is that MT lattice architecture may directly regulate signaling proteins. Consistent with recent studies (Meiring et al., 2025), GEF-H1 preferentially associated with compact lattice regions and we demonstrate here that it redistributes into short curved filaments following MT lattice expansion. Using curved filament formation as a cellular readout of lattice preference, we systematically compared multiple C1 domain-containing proteins (Choi et al., 2026) and found that lattice sensitivity is not a universal property of C1 domains. Whereas GEF-H1 displayed a strong preference for compact lattice regions, RASSF1A and RASSF1D preferentially associated with expanded lattice states despite containing closely related C1 domains. Moreover, whereas the isolated GEF-H1 C1 domain retained lattice sensitivity, the isolated RASSF1A and RASSF1D C1 domains failed to associate with MTs, indicating that lattice recognition depends not only on the C1 domain itself but also on cooperative interactions with additional regions of the protein or other MT-associated factors. These findings suggest that MT lattice state selectively organizes subsets of signaling proteins and may spatially regulate downstream signaling pathways through structural recognition of distinct lattice conformations.

Finally, we show that these principles extend to neurons, where many lattice-sensitive proteins are highly enriched and where MT organization is essential for intracellular transport, signaling, and mechanical stability. Tau and DCX in neurons exhibited responses to lattice modulation similar to those observed in non-neuronal cells, particularly within the somatic MT network, indicating that lattice-state-dependent protein organization is conserved in a physiologically specialized context. Interestingly, axonal tau association was comparatively resistant to Taxol-induced redistribution despite recent in situ cryo-EM studies demonstrating that axonal MTs adopt a stable GDP-bound expanded lattice state during neuronal differentiation (Zehr et al., 2026).

Together, these observations suggest that lattice spacing alone is insufficient to dictate protein recruitment in neurons and that additional neuron-specific features, including MAP composition, MT bundling, or higher-order cytoskeletal organization, modulate lattice accessibility and stabilize protein association within axons.

Together, our findings establish short, curved MT-associated filaments as the first in-cellulo readout of compact MT lattice preference and identify MT lattice spacing as a previously underappreciated mechanochemical regulatory layer that spatially organizes protein recruitment, signaling, and cytoskeletal architecture in cells. Future studies combining live-cell imaging with in situ cryo-electron tomography should reveal how nanoscale lattice remodeling gives rise to these curved domains and determine how mechanical forces reshape the MT lattice to regulate protein organization in both healthy and diseased states.

## ACKNOWLEDGEMENTS

N.M.R. acknowledges support from the Deutsche Forschungsgemeinschaft (DFG, German Research Foundation; project numbers 335549539 (GRK2381) and 587521045 (MO 5001/6-1), the DFG-funded microscope grant 434558229 (INST 37/1133-1 FUGG), and core funding from the Excellence Strategy at the University of Tübingen. J.M. was supported by the DFG Research Training Group GRK2381 (project number 335549539). This work was also supported by a DFG grant to S.P. (project number PY 121/2-1). We also thank Halil Kaan Türk (GTC neuroscience) and Melissa Lang Schnee (IFIB biochemistry) for their assistance with specific experiments. N.M.R. gratefully acknowledges many stimulating discussions with Sasha Bershadsky (Mechanobiology Institute, Singapore), which helped shape the conceptual framework of this study on microtubules as signaling hubs.

## AUTHOR CONTRIBUTIONS

N.M.R. conceptualized the project. J.M. designed and performed all experiments, with contributions from P.V. S.P. provided experimental assistance, reagents, and contributed to manuscript editing. P.V., L.T., and B.B. provided extensive experimental assistance, while B.B. also provided technical support and laboratory management. K.W. performed quantitative analysis of the MT structures. J.M. and N.M.R. wrote the manuscript. N.M.R. supervised and coordinated the project. All authors read and approved the final manuscript.

## METHOD

### Plasmids and constructs

All GEF-H1 (UniProt: Q92974), RASSF1A (UniProt: Q9NS23-2), and RASSF1D (UniProt: Q9NS23-1) constructs used in this study were generated from human codon-optimized sequences synthesized by GenScript using the CloneEZ cloning system. Full-length coding sequences were cloned into the N-terminal GFP expression vector pDEST-GFP- N1. C1-domain constructs corresponding to GEF-H1 (amino acids 28–100), RASSF1A (amino acids 1–110), and RASSF1D (amino acids 1–114) were similarly cloned into pDEST-GFP-N1 by GenScript. All other plasmids used in this study were obtained from Addgene and are listed in Supplementary Table 1.

### Cell Lines and iPSC culture

HelaM, COS-7 were grown in DMEM (Thermo Fisher Scientific) and RPE1 cells in RPMI (Thermo Fisher Scientific) supplemented with 10% FBS (Thermo Fisher Scientific) and 1% penicillin-streptomycin. Cells were kept at 37 °C with 5% CO_2_ in an enclosed incubator.

The following iPSC lines were obtained from the Jackson Laboratories (JAX) and iNDI consortium: KOLF2.1J(TO-NGN2, JIPSC002070). For the maintenance of iPSCs in culture, iPSCs were cultured on Geltrex (Life Technologies) coated dishes and maintained in Essential 8 Flex media (Thermo Fisher Scientific). The Rho-kinase (ROCK) inhibitor Y-27632 (EMD Millipore, 10 μM) was added to Essential 8 Flex media on the first day of plating and replaced with fresh media without ROCK inhibitor on the following day.

For i^3^neuronal differentiation, iPSCs were differentiated into cortical-like i^3^Neurons according to a previously described protocol based on the doxycycline inducible expression of Ngn2(Fernandopulle et al., 2018). A detailed protocol can be found at https://www.protocols.io/view/culturing-i3neurons-basic-protocol-6-n92ld3kbng5b/v1.

### DNA transfection

For transfection of COS-7 and HelaM cells, 1 μl Lipofectamine™ 2000 Transfection Reagent (Invitrogen) was used and for RPE1 cells, 3 μl Lipofectamine™ 2000 was used with the respective plasmids and visualized within 24-48 h. For transfection of i^3^Neuron, plasmids were transfected with 4 μl of Lipofectamine™ Stem Transfection Reagent (Invitrogen) on day 9 and visualized at least 72 h later. A detailed protocol can be found https://doi.org/10.17504/protocols.io.5qpvokx3bl4o/v1.

### Drugs

All drug treatments were performed on live cells. Unless otherwise stated, the same cells were imaged before and after drug treatment, allowing each cell to serve as its own internal control. Cells were first imaged under basal conditions, incubated with the indicated drug for at least 30 min or more, and then re-imaged. The compounds used in this study were Taxol (Sigma-Aldrich), Nocodazole (MedChemExpress), Laulimalide (Santa Cruz Biotechnology), and SiR-tubulin (Spirochrome), a fluorogenic MT probe that binds tubulin through a taxane-based mechanism. Taxol and Laulimalide were used at a final concentration of 10 μM and were diluted in complete culture medium for HeLaM and COS-7 cells or CM for i^3^Neurons. Nocodazole was used at a final concentration of 5 μM in complete culture medium. Unless otherwise stated, SiR-tubulin was used at a final concentration of 100 nM; where indicated, 500 nM SiR-tubulin was used to promote the formation of curved filament-like structures.

### Live imaging and fluorescent microscopy

For all imaging experiments, cells were seeded on glass-bottom ibidi dishes (ibidi GmbH) and transfected as described above. For live-cell imaging, COS-7, HeLaM and RPE1 cells were maintained in Live Cell Imaging Buffer (Life Technologies), whereas i^3^Neurons were maintained in CM. Imaging was performed in a stage-top incubation chamber maintained at 37 °C with a humidified atmosphere containing 5% CO₂. Live-cell imaging experiments were performed on the same cells before and after the indicated drug treatment. Fixed-cell imaging experiments were performed on independent samples. Images were acquired using a Zeiss LSM 980 confocal microscope equipped with Airyscan 2 using excitation wavelengths between 405 and 640 nm, as appropriate. Images were collected using a Plan-Apochromat 60× oil-immersion objective (NA 1.45).

### Immunofluorescence

For immunofluorescence staining of tyrosinated and detyrosinated microtubules in COS-7 and HeLaM cells, methanol fixation was used instead of paraformaldehyde (PFA) fixation, as PFA fixation caused tau and DCX to dissociate from microtubules and become predominantly cytoplasmic. Cells were fixed with pre-chilled methanol at −20 °C for 15 min, followed by three washes with PBS. Cells were then permeabilized with 0.1% Triton X-100 in PBS for 5 min and blocked for 1 h in 5% bovine serum albumin (BSA) in PBS. Primary antibodies were diluted in blocking buffer and incubated overnight (24 h) at 4 °C (see Supplementary Table 2 for antibody details and dilutions). After three PBS washes, cells were incubated with the appropriate secondary antibodies for 1 h at room temperature (Supplementary Table 2), followed by PBS washes before imaging.

For i^3^Neuron samples stained for tyrosinated, detyrosinated, acetylated, and βIII- tubulin, cells were fixed and extracted for 15 min in 4% (v/v) paraformaldehyde, 0.2% (v/v) glutaraldehyde, and 0.25% (v/v) Triton X-100 in CB at 37 °C, followed by two PBS washes. Free aldehyde groups were quenched by incubation with freshly prepared 1 mg/ml sodium borohydride in CB for 10 min, followed by three PBS washes. Cells were blocked for 30 min in 5% BSA in PBS and incubated overnight at 4 °C with primary antibodies (Supplementary Table 2). After three PBS washes, samples were incubated with Alexa Fluor-conjugated secondary antibodies (Thermo Fisher Scientific; Supplementary Table 2) for 1 h at room temperature, followed by three final PBS washes before imaging.

### Image analysis and quantification

Images were pseudocolor-coded, adjusted for brightness and contrast, cropped and/or rotated using the open-source image processing software FIJI (ImageJ)(Schindelin et al., 2012). Curved filament-like structures were quantified using the CurveTrace plugin for FIJI together with the accompanying analysis scripts(Katrukha et al., 2021). Images were exported as TIFF files and analyzed separately for DCX and tau. Because the exported TIFF files did not retain the original spatial calibration, the pixel size was manually restored before analysis (1 pixel = 0.042527 µm). Curvilinear DCX- and tau-positive structures were detected semi-automatically using CurveTrace. All detected traces were visually inspected and manually curated before being saved as regions of interest (ROIs). Each image was paired with its corresponding ROI file using matched filenames and analyzed using a custom FIJI macro. For each ROI, point coordinates were extracted and filament length was calculated as the cumulative Euclidean distance between consecutive points, followed by conversion from pixels to micrometers using the calibrated pixel size. Measurements were exported as CSV files and analyzed in Python using custom-written scripts. Length distributions were first examined to distinguish segmentation artifacts, punctate structures, and curved filaments. Structures ff0.2 µm were classified as segmentation-derived artifacts and excluded from further analysis.

Structures between 0.2 and 0.5 µm were classified as punctate structures and included only in the quantification of puncta. Structures ≥0.5 µm were classified as curved filaments and subjected to curvature analysis. For curvature measurements, each curved filament was fitted to a circle. The fitted radius was converted to micrometers, and the arc angle was calculated from the ratio of arc length to fitted radius. The root mean squared error (RMSE) between the traced coordinates and the fitted circle was used to assess the quality of the circular fit. Only curved filaments with an RMSE below the predefined threshold were retained for quantitative analysis, ensuring reliable estimation of filament length, radius, and arc angle.

### Statistical analysis

The methods for statistical analysis and sizes of the samples (n) are specified in the results section or figure legends for all quantitative data. Comparisons between two groups were performed using Student’s t-test or Welch’s t-test, whereas comparisons among three groups were performed using one-way ANOVA. Differences were accepted as significant for P ff 0.05. Prism version 10 (GraphPad Software) was used to plot, analyze and represent the data.

## Data availability

All data generated or analyzed during this study are included in this published article (and its Supplementary Material). Raw datasets generated during and/or analyzed during the current study are available from the corresponding author on request.

---

**Supplementary Figure 1.**
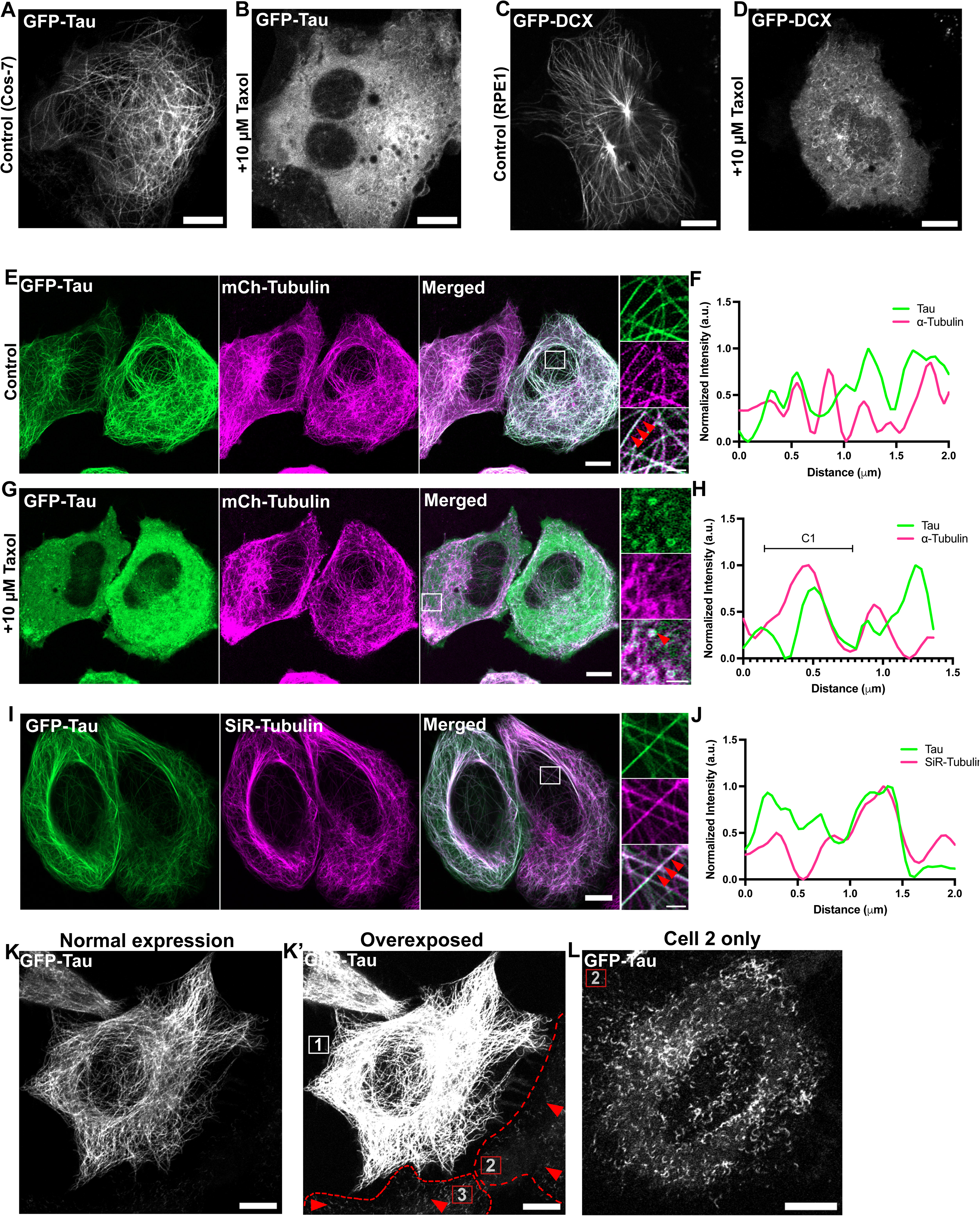
Taxol-induced curved filament formation is conserved across cell types and depends on tau expression levels. (A, B) Representative fluorescence images of COS-7 cells expressing EGFP–tau under control conditions (A) or following Taxol treatment (B). Similar to HeLaM cells, Taxol induces redistribution of tau into curved filament-like structures together with a diffuse cytoplasmic pool. Scale bars, 10 μm. (C, D) Representative fluorescence images of RPE1 cells expressing EGFP–DCX under control conditions (C) or following Taxol treatment (D). Similar to HeLaM cells, Taxol induces curved filament-like structures and cytoplasmic redistribution of DCX. Scale bars, 10 μm. (E–H) Representative fluorescence images of HeLaM cells co-expressing EGFP–tau (green) and mCherry– tubulin (magenta). Similar to SiR-tubulin labeling shown in Figure 1, EGFP–tau overlaps with mCherry–tubulin under control conditions (E, F) and redistributes into curved filament-like structures following Taxol treatment while remaining associated with an intact MT network (G, H). Scale bars, 10 μm; insets, 2 μm. (F, H) Line-scan intensity profiles of the regions indicated in (E) and (G), respectively. (I) Representative fluorescence image of a HeLaM cell expressing EGFP–tau (green) labeled with a low concentration of SiR-tubulin (100 nM; magenta). At low SiR-tubulin concentrations, tau remains uniformly associated with MTs, in contrast to the curved filament-like structures induced by higher SiR-tubulin concentrations (500 nM; Figure 1F). (J) Line-scan intensity profile of the region indicated in (I). (K, K’) Representative fluorescence images of HeLaM cells expressing different levels of EGFP–tau. Cells expressing higher levels of tau (Cell 1) display uniform MT decoration, whereas cells expressing lower levels of tau (Cells 2 and 3), visualized following brightness adjustment, exhibit spontaneous curved filament-like structures. (L) Higher-magnification view of Cell 2 shown in (K’), revealing numerous curved filament-like structures. These observations suggest that at low expression levels tau preferentially localizes to compact lattice regions, whereas at higher expression levels tau uniformly decorates and may compact MTs. Scale bars, 10 μm.

**Supplementary Figure 2.**
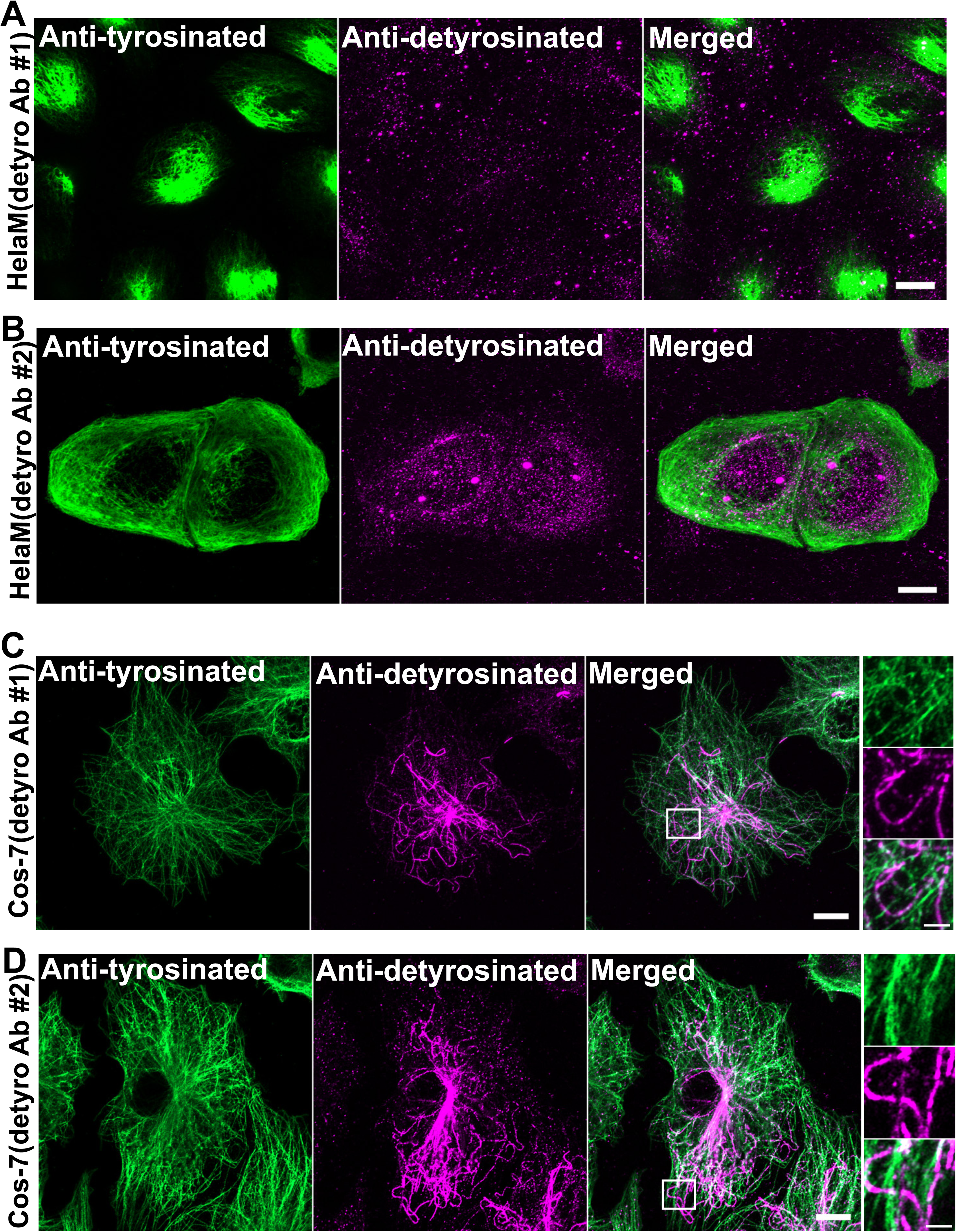
HeLaM cells lack detectable detyrosinated MTs. (A, B) Representative fluorescence images of HeLaM cells immunolabeled for tyrosinated tubulin (green) and detyrosinated tubulin (magenta) using two independent antibodies. No detectable detyrosinated MTs were observed with either antibody. Scale bars, 10 μm. (C, D) Representative fluorescence images of COS-7 cells immunolabeled for tyrosinated tubulin (green) and detyrosinated tubulin (magenta) using the same two antibodies. In contrast to HeLaM cells, COS-7 cells exhibited robust detyrosinated MT staining with both antibodies, confirming antibody specificity. Scale bars, 10 μm; insets, 2 μm.

**Supplementary Figure 3.**
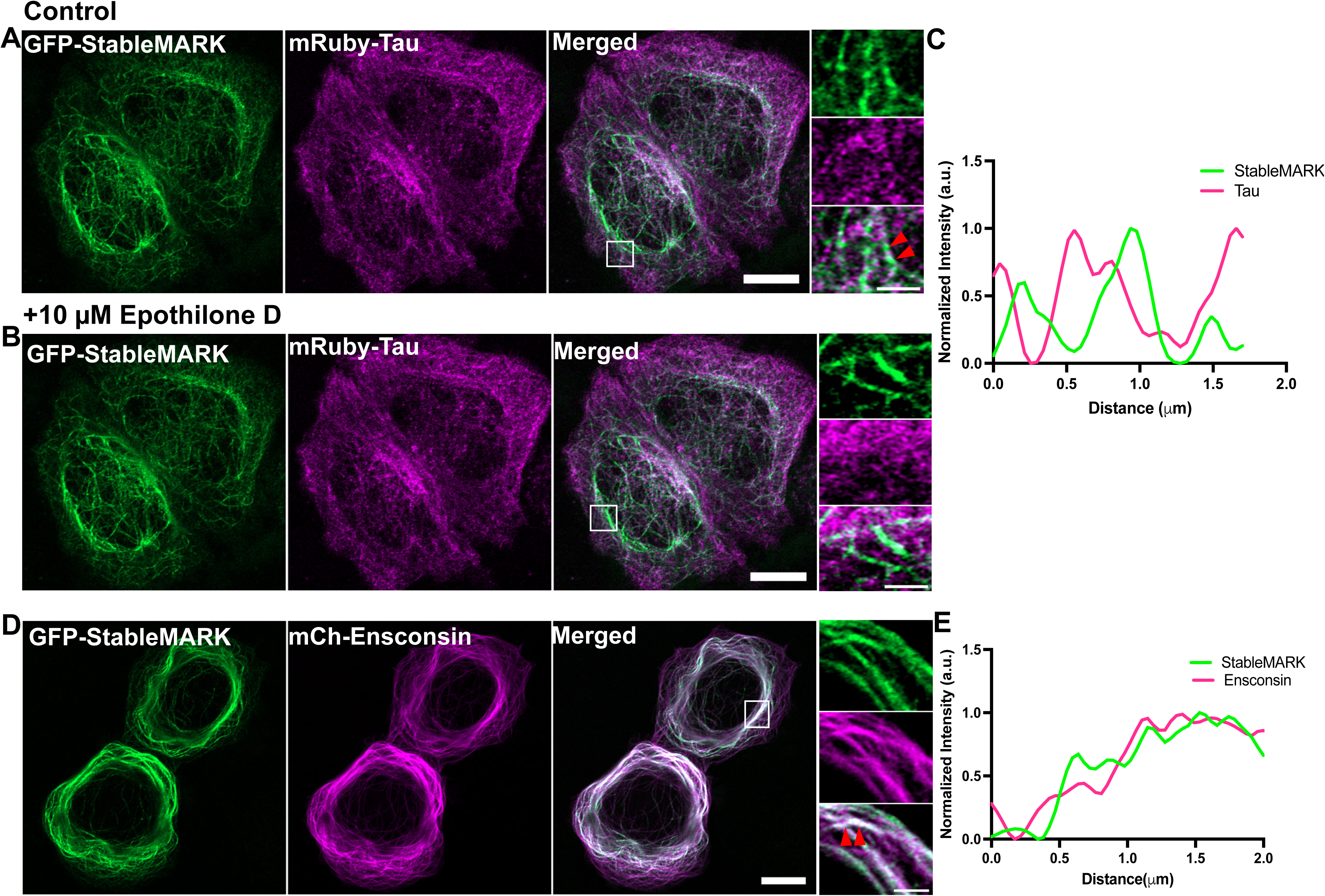
Expanded lattice–binding proteins remain associated with MTs following lattice expansion. (A, B) Representative fluorescence images of HeLaM cells co-expressing EGFP– StableMARK (green) and mRuby–tau (magenta) under control conditions (A) or following treatment with 10 μM epothilone D (B). Similar to Taxol (Figure 3), epothilone D induces dissociation of tau from MTs while StableMARK remains robustly associated with the MT network. Scale bars, 10 μm; insets, 2 μm. (C) Line-scan intensity profile of the region indicated in (A). (D) Representative fluorescence images of HeLaM cells co-expressing EGFP–StableMARK (green) and mCherry–ensconsin (magenta). Both proteins show extensive overlap along MTs, indicating preference for the same MT lattice state. Scale bars, 10 μm; insets, 2 μm. (E) Line-scan intensity profile of the region indicated in (D).

**Supplementary Figure 4.**
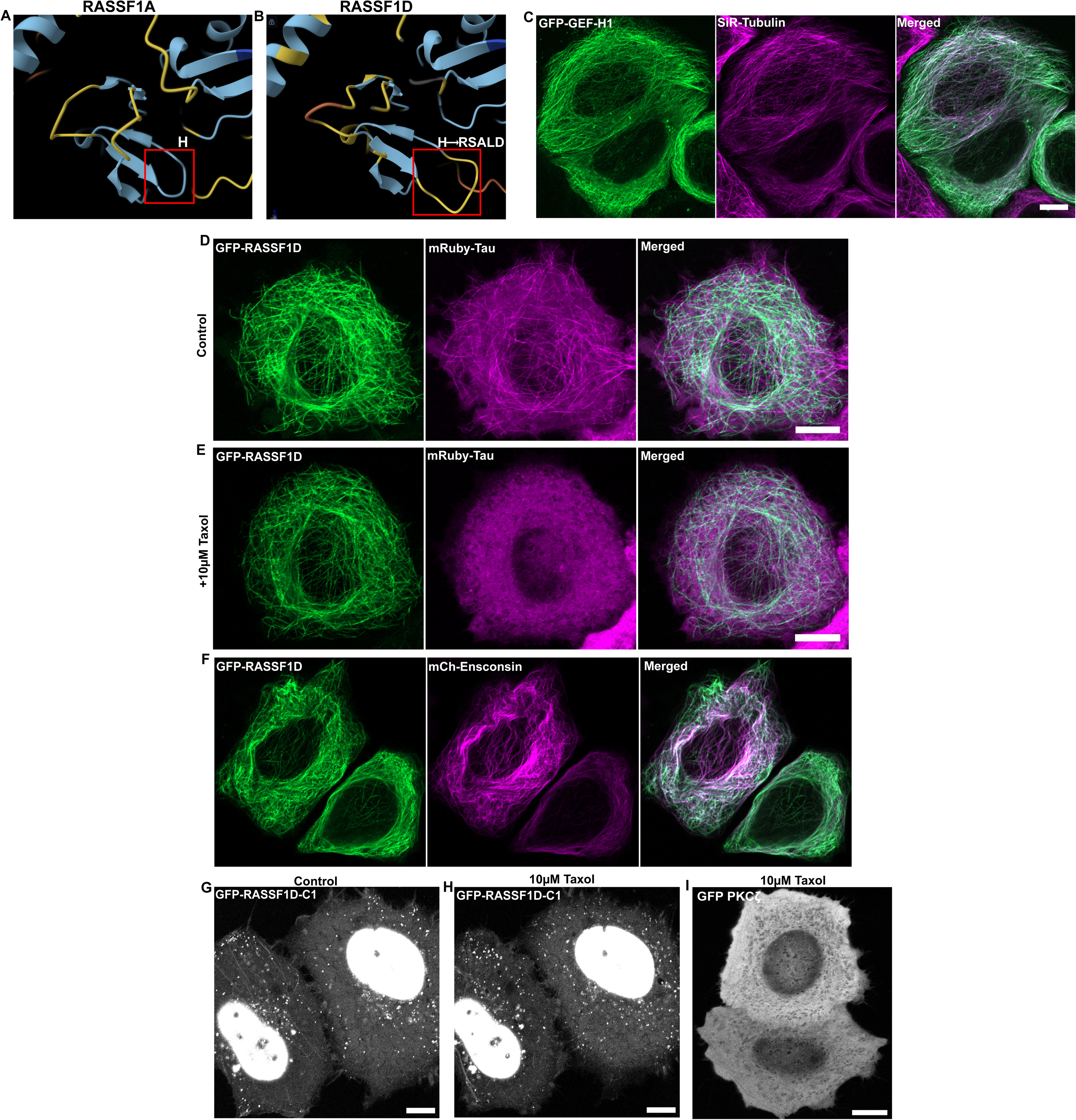
Structural comparison and lattice preference of C1 domain–containing signaling proteins. (A, B) AlphaFold-predicted structures of the C1 domains of human RASSF1A (A; UniProt Q9NS23-2) and RASSF1D (B; UniProt Q9NS23-1), highlighting the sequence difference within the C1 domain (H in RASSF1A versus RSALD in RASSF1D), which is predicted to generate a longer loop in the RASSF1D structure. (C) Representative fluorescence image of a HeLaM cell expressing EGFP–GEF-H1 (green) labeled with SiR-tubulin (100 nM; magenta). EGFP–GEF-H1 extensively overlaps with the MT network. Scale bars, 10 μm. (D, E) Representative fluorescence images of HeLaM cells expressing full-length EGFP–RASSF1D under control conditions (D) or following Taxol treatment (E). Similar to RASSF1A, RASSF1D remains associated with MT following Taxol treatment, whereas tau redistributes into a diffuse cytoplasmic pool. Scale bars, 10 μm. (F) Representative fluorescence image of a HeLaM cell co-expressing EGFP–RASSF1D (green) and mCherry–ensconsin (magenta). Both proteins extensively overlap, indicating preference for the same MT population. Scale bars, 10 μm. (G, H) Representative fluorescence images of HeLaM cells expressing the isolated EGFP–RASSF1D C1(aa 1-114) domain under control conditions (G) or following Taxol treatment (H). Similar to the isolated RASSF1A C1 domain, the RASSF1D C1 domain exhibits predominantly cytoplasmic and nuclear localization, lacks detectable MT association, and shows no redistribution following Taxol treatment. Scale bars, 10 μm. (I) Representative fluorescence images of the same HeLaM cell expressing EGFP–PKCζ shown in Figure 5E (before) and after Taxol treatment (I). Similar to control conditions, PKCζ remains predominantly cytoplasmic and shows no detectable MT association or redistribution following Taxol treatment. Scale bars, 10 μm.

**Supplementary Figure 5.**
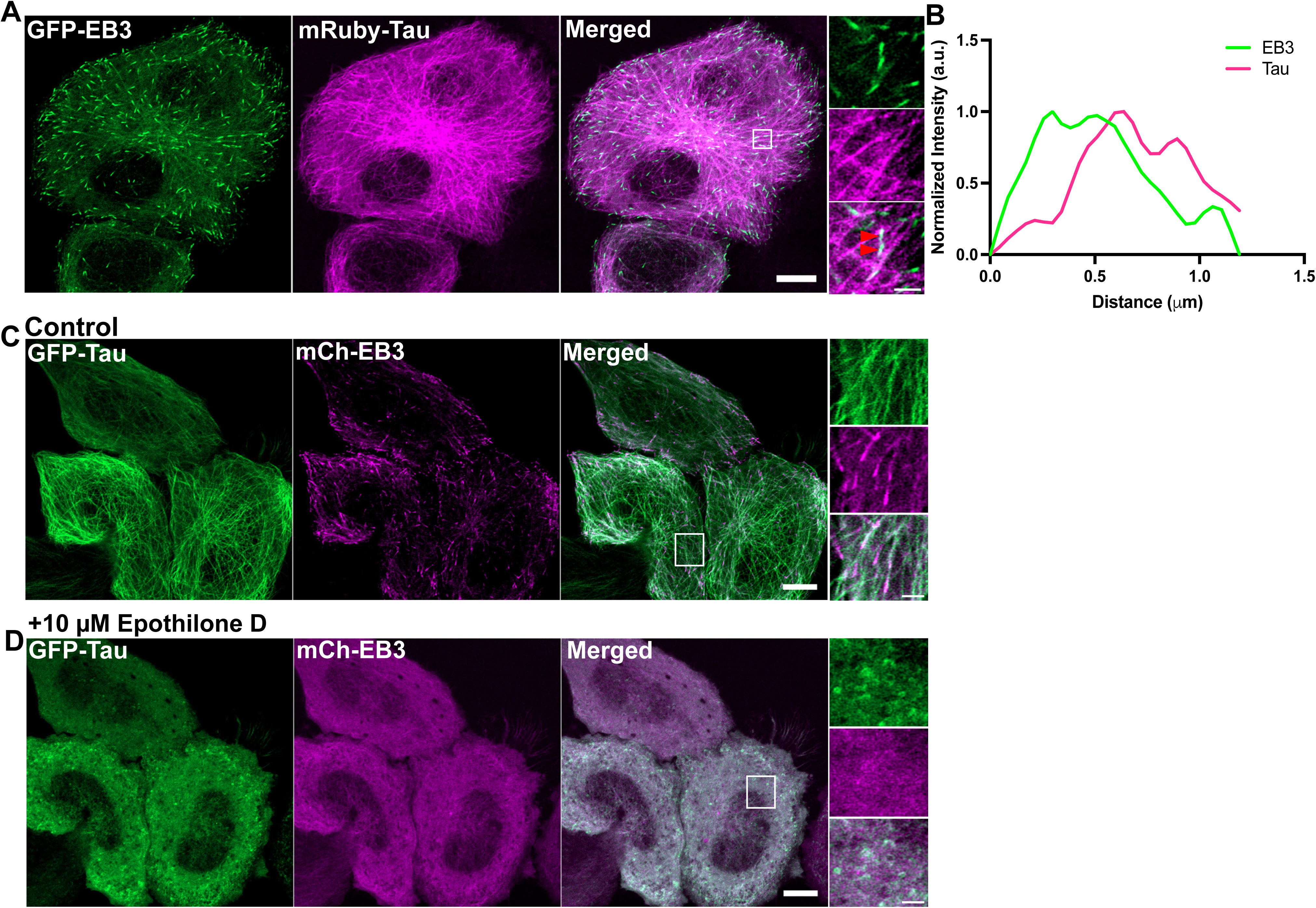
Supplementary Figure 5. EB3 and tau display distinct localization at MT plus ends and similar responses to epothilone. **D.** (A, B) Representative fluorescence images of HeLaM cells co-expressing EGFP–EB3 (green) and mRuby–tau (magenta). Line-scan analysis (B) shows that tau localizes immediately behind EB3-positive MT plus ends, with minimal overlap at the EB3-labeled tip. Scale bars, 10 μm; insets, 2 μm. (C, D) Representative fluorescence images of HeLaM cells co-expressing EGFP–Tau (green) and mCherry EB3 (magenta) under control conditions (C) or following treatment with 10 μM epothilone D (D). Similar to Taxol (Figure 6), epothilone D induces redistribution of both EB3 and tau from MTs into a diffuse cytoplasmic pool together with curved filament-like structures. Scale bars, 10 μm; insets, 2 μm.

**Supplementary Figure 6.**
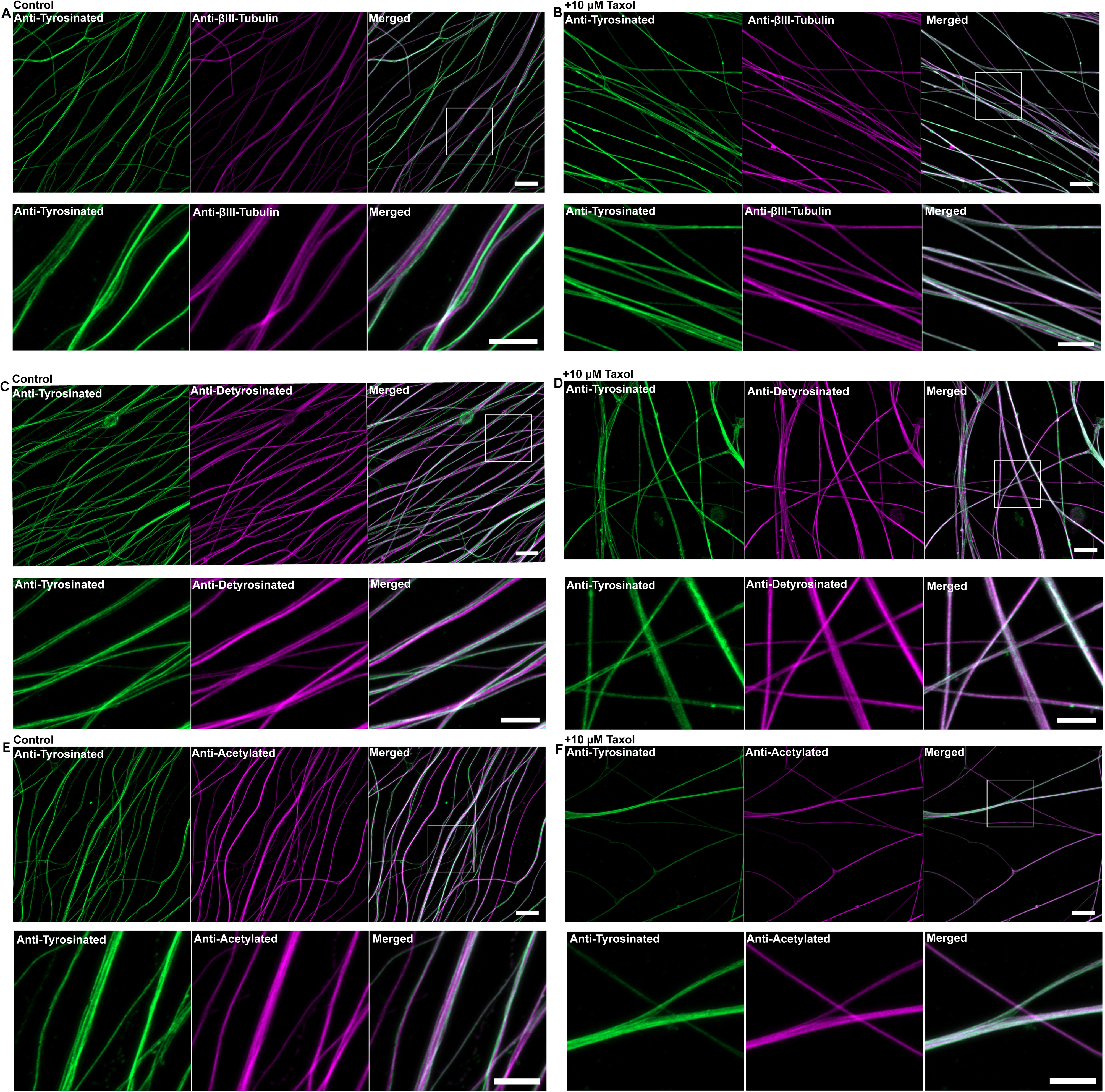
Acute Taxol treatment does not alter canonical tubulin PTMs in axonal and dendritic MTs. (A, B) Representative fluorescence images of axonal and dendritic regions of i^3^Neurons (days 18–22) immunolabeled for βIII-tubulin (magenta) and tyrosinated tubulin (green) under control conditions (A) or following Taxol treatment (B). Boxed regions are shown at higher magnification below each panel. Robust βIII-tubulin staining confirms the neuronal identity of the differentiated iPSC-derived cultures. Scale bars, 10 μm; insets, 5 μm. (C, D) Representative fluorescence images of axonal and dendritic regions of i^3^Neurons (days 18–22) immunolabeled for tyrosinated (green) and detyrosinated tubulin (magenta) tubulin under control conditions (C) or following Taxol treatment (D). Boxed regions are shown at higher magnification below each panel. Tyrosinated and detyrosinated MTs exhibit similar distributions under both conditions. Scale bars, 10 μm; insets, 5 μm. (E, F) Representative fluorescence images of axonal and dendritic regions of i^3^Neurons immunolabeled for tyrosinated (green) and acetylated tubulin (magenta) under control conditions (E) or following Taxol treatment (F). Acetylated MTs likewise show no obvious changes in distribution following Taxol treatment. Acetylated, tyrosinated, and detyrosinated tubulin were detected by immunofluorescence using specific antibodies. Scale bars, 10 μm; insets, 5 μm.

**Supplementary Video 1.** Live fluorescence imaging of a HeLaM cell expressing EGFP– tau (green) and labeled with high-concentration SiR-tubulin (500 nM, magenta), demonstrating overlap of tau-positive curved filaments with the underlying microtubule network. The movie is displayed at 5 frames/sec. Scale bar, 10 μm.

**Supplementary Video 2.** Live fluorescence imaging of a HeLaM cell co-expressing EGFP–DCX (green) and mCherry–tubulin (magenta) following Taxol treatment, showing the formation of curved filament-like and punctate DCX-positive structures that remain associated with the underlying microtubule network. Note the dynamic behavior of the DCX-positive curved filaments. The movie is displayed at 5 frames/sec. Scale bar, 10 μm.

**Supplementary Video 3.** Live fluorescence imaging of a HeLaM cell co-expressing EGFP–tau (green) and mCherry–ensconsin (magenta) following treatment with the MT lattice–expanding agent epothilone D. Tau redistributes into dynamic curved filament-like structures, whereas ensconsin remains filamentous and does not redistribute into curved filament-like structures. The movie is displayed at 5 frames/sec. Scale bar, 10 μm.

**Supplementary Video 4.** Live fluorescence imaging of a cropped region of a HeLaM cell co-expressing EGFP–GEF-H1 (green) and mCherry–ensconsin (magenta) following Taxol treatment. GEF-H1 redistributes into the cytoplasm and show dynamic curved filament-like structures, whereas ensconsin retains its filamentous distribution. The movie is displayed at 5 frames/sec. Scale bar, 2 μm.

**Supplementary Video 5.** Live fluorescence imaging of a HeLaM cell expressing the isolated EGFP–GEF-H1 C1 domain (green) following Taxol treatment. The GEF-H1 C1 domain redistributes into dynamic curved filament-like structures. The movie is displayed at 5 frames/sec. Scale bar, 10 μm.

**Supplementary Table 1.** List of antibodies used in this study.

| Target Protein | Company;<br>Catalog<br>number | Antibody<br>species | Working dilution for<br>immunofluorescence | Working<br>dilution for<br>immunoblotting |
| --- | --- | --- | --- | --- |
| βIII-Tubulin | Proteintech;<br>80713-1-RR | Rabbit | 1:500 | N/A |
| βIII-Tubulin | Proteintech;<br>66375-1-Ig | Mouse | 1:500 | N/A |
| Acetyl-alpha<br>Tubulin<br>(Lys40) | ThermoFisher;<br>322700 | Mouse | 1:500 | N/A |
| Detyrosinated-Alpha Tubulin | Abcam;<br>AB48389-1001 | Rabbit | 1:250 | N/A |
| Detyrosinated-Alpha Tubulin | RevMAb<br>Biosciences;<br>31-1335-00 | Rabbit | 1:500 | N/A |
| Tyrosinated-Alpha Tubulin<br>Clone YL1/2 | Sigma Aldrich;<br>MAB1864-I | Rat | 1:500 | N/A |
| **Secondary Antibodies** |  |  |  |  |
| Alexa Flour™<br>488 donkey-<br>anti-rabbit | ThermoFisher;<br>3148296 | Donkey | 1:500 | N/A |
| Alexa Flour™<br>488 donkey-<br>anti-mouse | ThermoFisher;<br>3281113 | Donkey | 1:500 | N/A |
| Alexa Flour™<br>555 goat-<br>anti-rabbit | ThermoFisher;<br>3148228 | Goat | 1:500 | N/A |
| Alexa Flour™<br>555 goat-<br>anti-mouse | ThermoFisher;<br>3174391 | Goat | 1:500 | N/A |
| Alexa Flour™<br>647 donkey-<br>anti-rabbit | ThermoFisher;<br>3219281 | Donkey | 1:500 | N/A |
| Alexa Flour™<br>647 donkey-<br>anti-mouse | ThermoFisher;<br>3141854 | Donkey | 1:500 | N/A |

**Supplementary Table 2.** List of plasmids used in this study.

| Plasmid | Company; Catalog number |
| --- | --- |
| EGFP-Tau | Addgene;<br>46904 |
| mRuby2-MAPTau-C-10 | Addgene;<br>55904 |
| EGFP-DCX | Addgene;<br>32852 |
| mCherry-Tubulin | Gift from Alexander Bershadsky (MBI,<br>NUS) |
| GFP-Ensconsin | Addgene;<br>26741 |
| mCherry-Ensconsin | Addgene;<br>26742 |
| EGFP-KIF5B-Rigor | Addgene;<br>172204 |
| EGFP-EB3 | Gift from Alexander Bershadsky (MBI,<br>NUS) |
| mCherry-EB3 | Gift from Alexander Bershadsky (MBI,<br>NUS) |
| GFP-MAP1B | Addgene;<br>44396 |
| mApple-MAP4-C-10 | Addgene;<br>54923 |
| mEGFP-TPX2-N-10 | Addgene;<br>56298 |
| EGFP-PKCZ | Addgene;<br>110512 |
| pDEST53-Flag(3x)-GFP-6X-Histag-GEF-H1 | Genscript |
| pDEST53-Flag(3x)-GFP-6X-Histag-GEF-H1-C1 | Genscript |
| pDEST53-Flag(3x)-<br>mRuby3-6X-Histag-GEF-H1 | Genscript |
| pDEST53-Flag(3x)-GFP-6X-Histag-RASSF1A | Genscript |
| pDEST53-Flag(3x)-GFP-6X-Histag-RASSF1A-C1 | Genscript |
| pDEST53-Flag(3x)-GFP-6X-Histag-RASSF1D | Genscript |
| pDEST53-Flag(3x)-GFP-6X-Histag-RASSF1D-C1 | Genscript |

